# Blocking gephyrin phosphorylation or microglia BDNF signaling prevents synapse loss and reduces infarct volume after ischemia

**DOI:** 10.1101/2020.04.22.055087

**Authors:** Teresa Cramer, Raminder Gill, Zahra S Thirouin, Markus Vaas, Suchita Sampath, Fanny Martineau, Sara B. Noya, David Colameo, Philip K.-Y. Chang, Peiyou Wu, Philip A Barker, Steven A. Brown, Rosa C. Paolicelli, Jan Klohs, R. Anne McKinney, Shiva K. Tyagarajan

**Author notes:** equal contribution (Teresa Cramer, Raminder Gill, Zahra S Thirouin). **Co-senior:** and (R. Anne McKinney, Shiva K. Tyagarajan).

## Abstract

Microglia interact with neurons to facilitate synapse plasticity; however, signal transducers between microglia and neuron remain unknown. Here, using *in vitro* organotypic hippocampal slice cultures and transient MCAO in genetically-engineered mice *in vivo*, we report that at 24 h post-ischemia microglia release BDNF to downregulate glutamatergic and GABAergic synapses within the peri-infarct area. Analysis of the CA1 hippocampal formation *in vitro* shows that proBDNF and mBDNF downregulate glutamatergic dendritic spines and gephyrin scaffold stability through p75^NTR^ and TrkB receptors respectively. Post-MCAO, we report that in the peri-infarct area and in the corresponding contralateral hemisphere similar neuroplasticity occur through microglia activation and gephyrin phosphorylation at Ser268, Ser270 *in vivo*. Targeted deletion of the *Bdnf* gene in microglia or *Gphn*S268A/S270A (phospho-null) point-mutations protect against ischemic brain damage, neuroinflamation and synapse downregulation normally seen post-MCAO. Collectively, we report that gephyrin phosphorylation and microglia derived BDNF faciliate synapse plasticity after transient ischemia.

## Introduction

Ischemic stroke is a leading cause of death and long-term disability worldwide. The annual mortality rate of 5.5 million is further compounded by high morbidity as up to 50% survivors are chronically disabled ^1^.Therapeutic approaches to CNS ischemia developed in the laboratory have focused on mechanisms contributing to ischemic damage, namely excitotoxicity, oxidative stress and inflammation ^2, 3^. Unfortunately to date clinical trails targeting glutamate receptors ^4^, GABA receptors ^5^, calcium channels ^6^, sodium channels ^7^and free radicals ^8^have all failed. The lack of treatment options is directly related to our poor understanding of the possible mechanisms underlying the disease. As the brain has developed inherent mechanism(s) for self-preservation; gaining insights into these protective measures may thus provide a way forward to counteract ischemic brain injury.

Typically, rapid cell death occurs in the ischemic core but intriguingly neurons in the peri-infarct area, a region with constrained blood flow and partially preserves energy metabolism, survive. Within the peri infarct the propagating neuronal depolarization in combination with impairment of glia function causes an increased extracellular concentration of ions and neurotransmitters resulting in neuronal functional perturbations ^9^. This is accompanied by reductions in both excitatory dendritic spines ^10^ and GABAergic synapses ^11^. Interestingly, the neurotrophin brain-derived neurotrophic factor (BDNF) can decrease cell death and ischemic core volume leading to improvement of neurological outcome after experimental stroke either upon overexpression *in vivo* using genetic methods ^12^ or upon exogenous application ^13^. Correspondingly, inhibition of BDNF exacerbates ischemic damage ^14^.

Under physiological conditions, glutamatergic neurotransmission induces BDNF expression ^15^. BDNF is expressed as a proprotein, proBDNF, and is subsequently processed to its mature form mBDNF ^16^. proBDNF preferentially binds to the low-affinity nerve growth factor receptor p75^NTR^ and negatively regulates dendritic spine stability through Rho GTPases, Rho/Rac1 activation ^17^. mBDNF signals through the neurotrophin receptor tyrosine kinase B (TrkB) to enhance excitatory neurotransmission ^18^. mBDNF can also bind to p75^NTR^ receptors, albeit with much lower affinity ^19^. At inhibitory GABAergic synapses, mBDNF induces the internalization of GABA_A_ receptors ^20^ and downregulation of the main inhibitory synapse scaffolding protein gephyrin ^21^, thereby reducing GABAergic transmission in principal cells.

Ischemia also activates inflammatory pathways that subsequently recruit leukocytes to the injured area of the brain ^22^. Immune cells contribute to both neuroprotection and programmed cell death ^23^, suggesting that the temporal window of inflammation might determine cell survival or death. The immune response signalling events must be counteracted to mitigate tissue damage and re-establish homeostasis. Microglia the resident immune cells in the central nervous system (CNS) are the primary responders during defense. They clear cellular debris as part of the tissue repair and wound healing processes ^24, 25^. In recent years, microglia have also been shown to play an essential role in synapse pruning during postnatal brain development ^26^. Microglia activation can be triggered by acute insult, causing process elongation and increased expression of marker proteins like IBA1 and CD11b ^27^. Microglial activation can protect the brain, albeit the precise cellular and molecular mechanisms for microglia influenced neuroprotection remain unclear. Microglia processes can directly sense synaptic activity ^28^, and can regulate neuronal calcium load and functional connectivity through neuronal mitochondrial function and P2Y12 receptor activation on contacting microglia ^29^. Subsequent inflammation induced by lipopolysaccharide (LPS) activates downstream calcium-calmodulin-dependent kinase (CaMKIV), cyclic AMP response element binding protein (CREB) phosphorylation and BDNF protein increase facilitate neuron survival after cortical injury ^30^.

It has been reported that BDNF administered either intravenously ^13^, with viral vectors ^12^, or by addition of the bioactive high-affinity TrkB agonist, 7,8-dihydroxyflavone, can protect neurons from apoptosis and decrease infarct volumes in animal models of stroke ^31^. These findings suggest that elevated BDNF is beneficial for recovery after stroke. However, the mechanisms underlying the beneficial effect of BDNF post-ischemia remain unclear. Here, we set out to assess the physiological mechanism(s) that are triggered upon ischemic brain damage to enable tissue repair and neural network reorganization. Using organotypic hippocampal slice cultures and the oxygen-glucose deprivation (OGD) cellular model of ischemia, we report that within the first 90 min post-ischemia, proBDNF via p75^NTR^ disrupts glutamatergic, and mBDNF via TrkB disrupts GABAergic neurotransmission. We found that ERK1/2 and GSK3β pathways downstream of TrkB phosphorylate gephyrin at Ser 268 and Ser 270 residues resulting in GABAergic synapse loss. Using transient middle cerebral artery occlusion (MCAO) in wildtype and genetically-engineered mice, we uncover a central role for BDNF derived from microglia in influencing gephyrin phosphorylation downstream of TrkB receptor signaling. Using pharmacological depletion of microglia, CRISPR/Cas9 generated *Gphn*S268A/S270A mutant mouse, and *Bdnf* gene deletion from microglia, we consistently demonstrate reduced microglial activation and enhanced synapse preservation at 24 h post MCAO. Collectively, these observations unravel microglia derived BDNF as the signal transducer linking microglia and neurons, to activate ERK1/2 and GSK3 β pathways to influence glutamatergic and GABAergic synapse integrity through gephyrin phosphorylation.

## Results

### OGD causes glutamatergic and GABAergic synapse downregulation

To understand the mechanisms of BDNF action in ischemia we started with an *in vitro* model of ischemia, OGD, in organotypic hippocampal slice cultures obtained from transgenic mice that express myristoylated GFP in a subset of CA1 pyramidal neurons ^32^ and studied synaptic changes in the CA1 area after OGD (4 min) and recovery at 90 min and 24 h. First, we confirmed that we had induced hypoxia with OGD by measuring hypoxia-inducible factor 1 α (HIF1α) expression ^33^ and found 1.5-fold increase in HIF1α expression in area CA1 90 min after OGD (Suppl. Fig. 1a, b). We then determined glutamatergic synapse alterations after OGD by measuring changes in dendritic spines ^34^. We observed an overall significant reduction in spine density on CA1 pyramidal neurons at both 90 min and 24 h following OGD compared to control cultures (Fig. 1a, b; Supp. Fig. 1b). The mushroom and long-thin subtype of spines were particularly affected (Fig. 1b).

**Figure 1.**
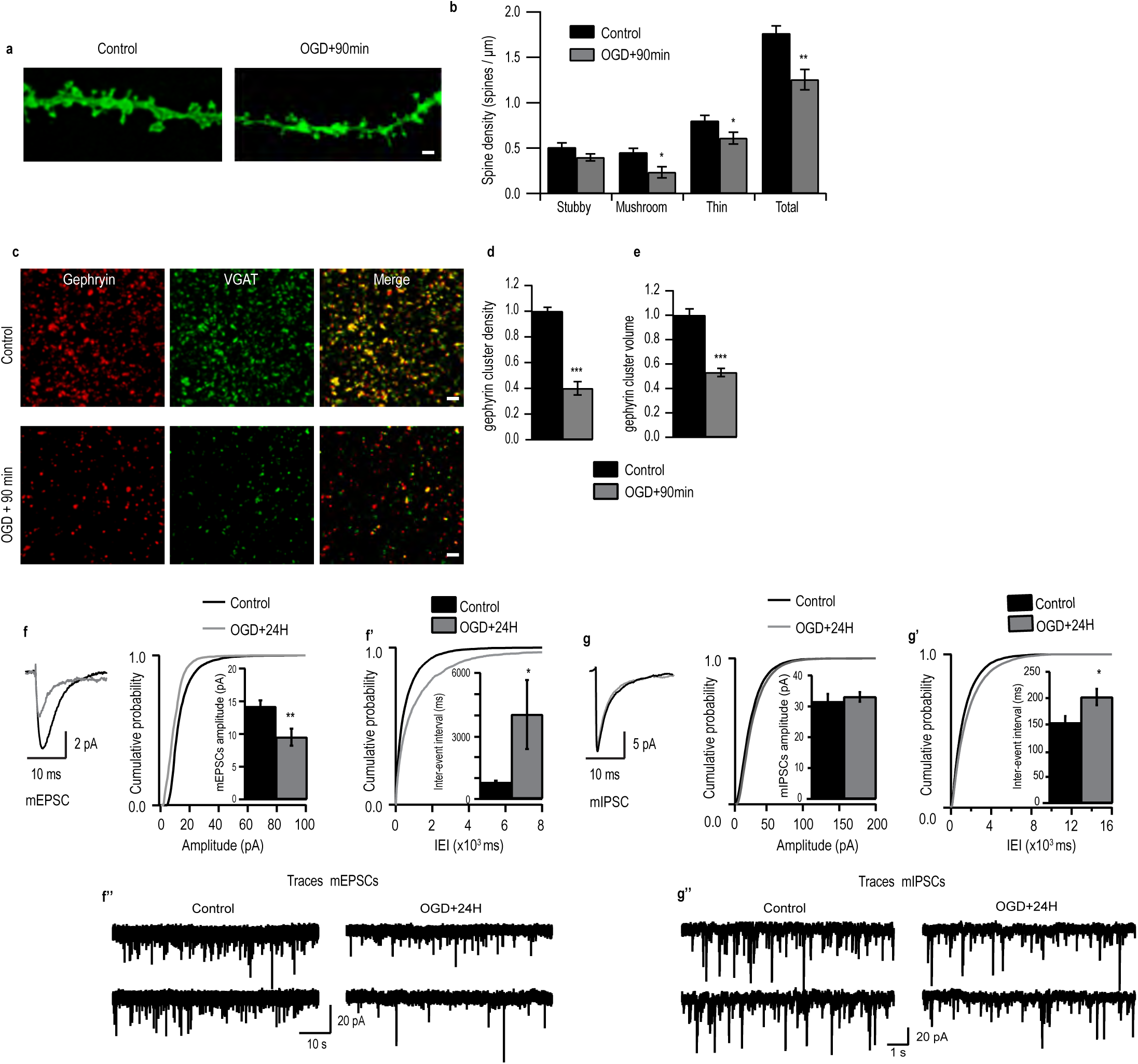
OGD induces morphological and functional deficits in excitatory synapses. **(a)** Example tertiary dendrites from CA1 pyramidal neurons expressing myristoylated-eGFP in control versus 90 min post-OGD organotypic hippocampal cultures. Scale = 2 µm. **(b)** Quantification of dendritic spines categorized into stubby, mushroom and long thin subtypes (\**p* < 0.05 and \*\**p* < 0.01, two-tailed independent Student’s *t*-test). Total spine density (spines/µm of dendrite): Control – 1.76 ± 0.08 (*n* = 8); OGD – 1.25 ± 0.11 (*n* = 8). **(c)** Example images of maximum intensity projections of area CA1 immunostained for gephyrin and VGAT in control cultures versus 90 min following OGD. **(d)** Quantification of number of gephyrin clusters per confocal stack (consisting of five 512×512 pixel z-planes each; \*\*\**p* < 0.0001, two-tailed independent Student’s *t*-test; gephyrin cluster values were normalized to control). Mean number (A.U.): Control – 1.00 ± 0.03 (*n* = 10 slices); OGD – 0.40 ± 0.05 (*n* = 10 slices). **(e)** Quantification of the total volume of gephyrin cluster (\*\*\**p* < 0.0001, two-tailed independent Student’s *t*-test). Mean volume (A.U.): Control – 1.00 ± 0.05; OGD – 0.53 ± 0.03. Data shown as mean ± s.d. (n=30 cells). **(f)** Cumulative probability histogram and mean ±SEM amplitude (*p* < 0.001, Kolmogorov-Smirnov test). **(f’)** Cumulative probability histogram and mean ±SEM for IEIs of mEPSCs (*p* < 0.05, Kolmogorov-Smirnov test). **(f’’)** sample mEPSC traces from control cells and OGD cells. **(g)** Cumulative probability histogram and mean ±SEM amplitude of mIPSCs (*p* < 0.001, Kolmogorov-Smirnov test). **(g’)** Cumulative probability histogram and mean ±SEM for IEIs of mIPSCs (*p* < 0.05, Kolmogorov-Smirnov test). **(f’’)** sample mEPSC traces from control cells and OGD cells. Data shown as mean ± s.d.

Next, we evaluated OGD-induced changes at GABAergic synapses in area CA1 at 90 min and 24 h following OGD. We immunolabeled for inhibitory presynaptic VGAT and postsynaptic inhibitory scaffolding protein gephyrin. We found a significant overall downregulation in gephyrin cluster density 90 min following OGD compared to control (Fig. 1c-e) in the *Stratum Radiatum*. However, after 24 h gephyrin cluster density remained significantly reduced, while cluster volume had recovered to baseline (Suppl. Fig. 1b’-b’’) similar to untreated cells.

To determine whether these morphological changes were accompanied by a functional deficit, we recorded AMPA-mediated miniature excitatory postsynaptic currents (mEPSCs) from CA1 pyramidal neurons within 24 h after OGD (Fig. 1f-f’). We found that the input resistance and resting membrane potential of CA1 hippocampal pyramidal cells in OGD and sister untreated cells were similar, suggesting that OGD does not impact receptor open probability or intracellular chloride concentration. However, we found a significant decrease in the mEPSC amplitude in OGD-treated slices compared to control (Fig. 1f). Similarly, the inter-event-interval (IEI) of mEPSC of OGD cells were increased compared to control (Fig. 1f’). Taken together, dendritic spine loss is mirrored by functional loss of excitatory synapses 24 h after OGD. Subsequently, to determine whether inhibitory circuitry was also affected we recorded GABA_A_-mediated miniature inhibitory postsynaptic current (mIPSC) within 24 h of OGD induction. mIPSC analysis showed no observable changes in the amplitude of mIPSC after OGD compared to control slices (Fig. 1g). However, a significant increase in the IEI in OGD-treated slices was observed (Fig. 1g’). These functional data recapitulate the morphological observations that inhibitory synapse loss after OGD does not recover, but total GABA_A_Rs at synaptic sites within the existing synapses recover at 24 h after OGD.

### Scavenging BDNF after OGD using TrkB-Fc rescues OGD-induced synapse deficit

As BDNF is upregulated after ischemia ^35, 36^, we assessed whether BDNF signaling contributed to synapse loss on CA1 pyramidal neurons after OGD. We scavenged proBDNF and mBDNF using chimeric TrkB-Fc (10μg/mL) and exposed organotypic hippocampal slices to 4 min OGD. Dendritic spine quantification in CA1 pyramidal neurons showed prevention of total spine density loss caused by OGD in TrkB-Fc treated cultures in comparison to untreated cultures (Fig. 2a, b). Specifically, the OGD-induced decrease in mushroom and long-thin subtype of dendritic spines was prevented by TrkB-Fc (Fig. 2a, b). We could confirm that TrkB-Fc scavenges both pro- and mBDNF by performing co-immunoprecipitation against TrkB-Fc and Western blot analysis against proBDNF and mBDNF (Fig. 2b’). Once we confirmed that TrkB-Fc also scavenges mBDNF, we assessed whether TrkB-Fc treatment also protects GABAergic synapses. We found that post-synaptic gephyrin clustering was protected by TrkB-Fc in OGD treated cultures (Fig. 2c-e). Importantly, TrkB-Fc caused no detectable effect in control cultures (not exposed to OGD).

**Figure 2.**
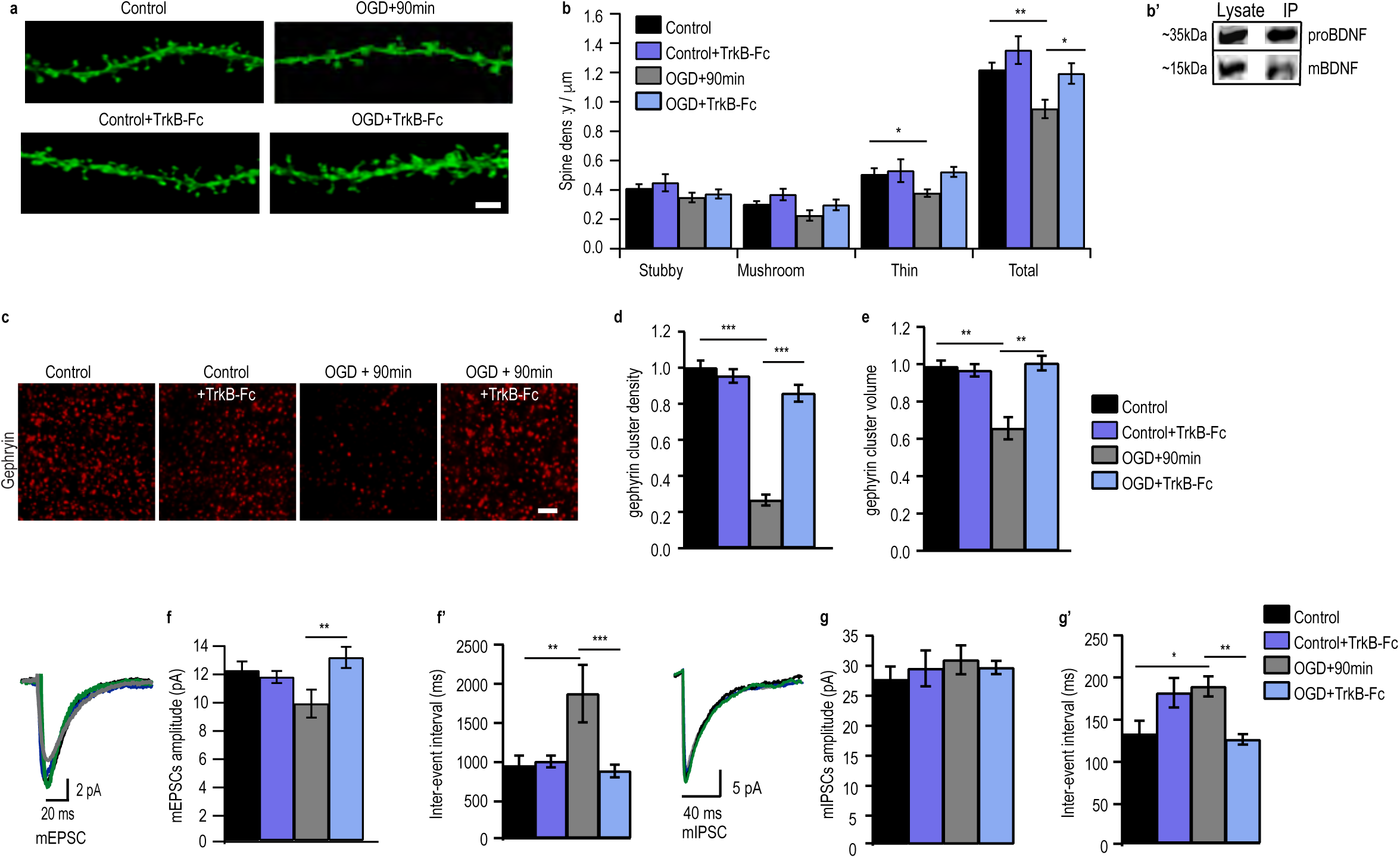
Scavenging BDNF rescues OGD-induced synapse deficits. **(a)** Example dendrites from CA1 neurons expressing myristoylated-eGFP with and without TrkB-fc treatment. Scale = 2 µm. **(b)** Quantification of dendritic spines (\**p* < 0.05, Two-way ANOVA with Bonferroni multiple comparison test). Total spine density (spines/µm of dendrite): Control – 1.22 ± 0.05 (*n* = 16); Control+TrkB-Fc – 1.40 ± 0.10 (*n* = 12); OGD – 0.95 ± 0.06 (*n* = 12); OGD+TrkB-Fc – 1.19 ± 0.07 (*n* = 14). **(b’)** Immunoprecipitation using TrkB-Fc chimera and WB against proBDNF or mBDNF. **(c)** Example images of maximum intensity projections of organotypic hippocampal slices immunostained for gephyrin with and without TrkB-Fc treatment. Scale = 2 µm. **(d)** Quantification of number of gephyrin puncta per confocal stack (consisting of five 512×512 pixel z-planes each; \*\*\**p* < 0.0001, Two-way ANOVA with Bonferroni multiple comparison test; all gephyrin values were normalized to control). Mean number (A.U.): Control – 1.00 ± 0.04 (*n* = 15); Control+TrkB-Fc – 0.95 ± 0.04 (*n* = 9 slices); OGD – 0.27 ± 0.03 (*n* = 13); OGD+TrkB-Fc – 0.86 ± 0.05 (*n* = 13). **(e)** Quantification of gephyrin puncta volume (\*\*\**p* < 0.0001, Two-way ANOVA with Bonferroni multiple comparison test). Mean volume (A.U.): Control – 1.00 ± 0.03; Control+TrkB-Fc – 0.98 ± 0.03; OGD – 0.67 ± 0.06; OGD+TrkB-Fc – 1.02 ± 0.04. Data shown as mean ± s.d. (n=30 cells). **(f)** Cumulative probability histogram of mean amplitude (*p* < 0.001, Two-way ANOVA with Bonferroni multiple comparison test). **(f’)** Cumulative probability histogram for IEIs of mEPSCs (*p* < 0.05, Two-way ANOVA with Bonferroni multiple comparison test). **(g)** Cumulative probability histogram of mean amplitude of mIPSCs (*p* > 0.05, Two-way ANOVA with Bonferroni multiple comparison test). **(g’)** Cumulative probability histogram for IEIs of mIPSCs (*p* < 0.05, Two-way ANOVA with Bonferroni multiple comparison test). Data shown as mean ± s.d.

To determine whether morphological synapse protection is recapitulated functionally, we recorded excitatory AMPA-mediated mEPSC from CA1 pyramidal neurons from all groups. We found that the OGD-induced reduction of mEPSC amplitudes 24 h following OGD was prevented by TrkB-Fc treatment, being comparable to treated and untreated controls (Fig. 2f; Suppl. Fig. 1c-c’’). Similarly, increase in IEI was also prevented by TrkB-Fc treatment as seen 24 h after the induction of OGD, with values similar to treated and untreated control slices (Fig. 2f’; Suppl. Fig. 1c’), indicating that the reduced occurrence of mEPSC after OGD was due to BDNF signaling.

To test if TrkB-Fc also prevented changes in inhibitory transmission we recorded GABA_A_- mediated mIPSC from CA1 pyramidal neurons from all groups. We found that mIPSC amplitudes were comparable in all groups (Fig. 2g, g’; Suppl. Fig. 1d). The previously observed increase in IEI caused by OGD was prevented with TrkB-Fc treatment to control levels (Fig. 2g’; Suppl. Fig. 1d’), confirming that the OGD-induced decrease in glutamatergic and GABAergic synapse loss is mediated by BDNF.

### proBDNF and mBDNF signal via p75^NTR^ and TrkB receptors to induce glutamatergic and GABAergic synapse loss respectively after ischemia

Next we investigated the molecular pathways involving BDNF-mediated synapse loss at 90 min post OGD in organotypic hippocampal slice cultures. To specifically investigate the contribution of proBDNF on OGD-induced dendritic spine loss, we used blocking antibodies to either inhibit proBDNF or p75^NTR^. Pre-treatment of OGD slices with either anti-proBDNF or anti-p75^NTR^ antibodies prevented OGD-induced dendritic spine loss (Fig. 3a-c; P=0.37). Additionally, treatment of OGD slices with anti-proBDNF or anti-p75^NTR^ antibody did not prevent gephyrin cluster loss after OGD (Fig. 3d; P=0.52). These findings indicate that pro-BDNF signaling through p75^NTR^ to specifically induces excitatory synapses loss following OGD.

**Figure 3.**
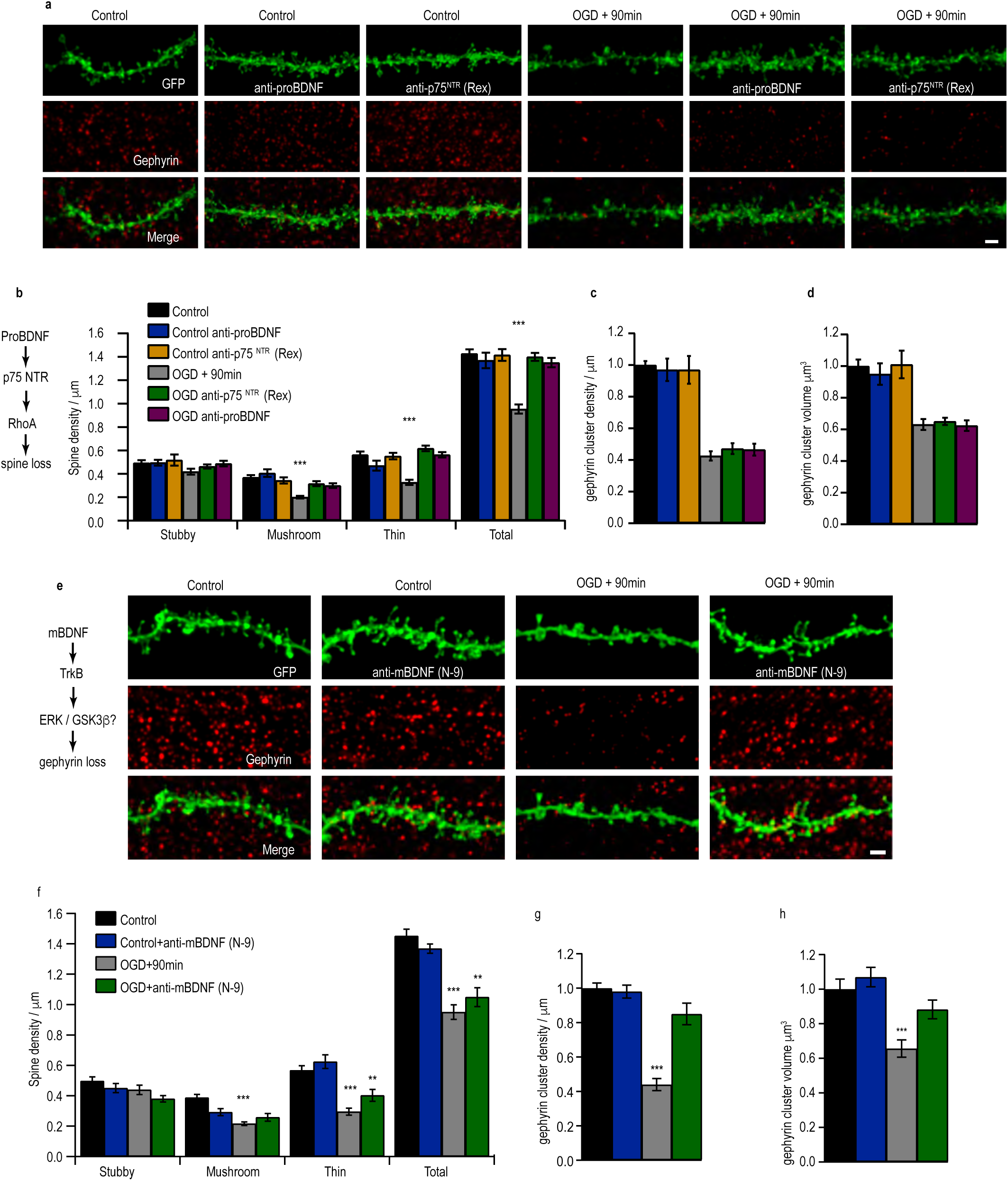
Differential proBDNF and mBDNF signaling induce synapse loss after ischemia. **(a)** Example tertiary dendrites from CA1 pyramidal neurons of organotypic hippocampal slices immunostained for gephyrin, with or without anti-proBDNF or anti-p75^NTR^ (Rex) treatment before OGD. Scale = 2 µm. **(b)** Dendritic spine quantification (\*\*\**p* < 0.0001 Two-way ANOVA with Bonferroni multiple comparison test). Total spine density (spines/µm of dendrite): Control – 1.43 ± 0.04 (*n* = 15); Control+anti-proBDNF – 1.37 ± 0.07 (*n* = 15); Control+anti-p75 ^NTR^ – 1.42 ± 0.05 (*n* = 15); OGD – 0.95 ± 0.04 (*n* = 39); OGD+anti-proBDNF – 1.40 ± 0.03 (*n* = 30); OGD+anti-p75^NTR^ – 1.35 ± 0.04 (*n* = 32). **(c)** Quantification of number of gephyrin cluster density (consisting of five 512×512 pixel z-planes each; *p* = 0.28, Two-way ANOVA with Bonferroni multiple comparison test, compared to control and treated controls; all gephyrin values were normalized to control). Mean number (A.U.): Control – 1.00 ± 0.03 (*n* = 18 slices); Control+anti-proBDNF – 0.97 ± 0.07 (*n* = 6); Control+anti-p75^NTR^ – 0.97 ± 0.06 (*n* = 6); OGD – 0.43 ± 0.03 (*n* = 20 slices); OGD+anti-proBDNF – 0.47 ± 0.03 (*n* = 16 slices); OGD+anti-p75^NTR^ – 0.47 ± 0.04 (*n* = 9). **(d)** Quantification of gephyrin puncta volume (*p* = 0.57, Two-way ANOVA with Bonferroni multiple comparison test, compared to control and treated controls). Mean volume (A.U.): Control – 1.00 ± 0.04; Control+anti-proBDNF – 0.95 ± 0.07; Control+anti-p75^NTR^ – 1.01 ± 0.09; OGD – 0.63 ± 0.03; OGD+anti-proBDNF – 0.65 ± 0.02; OGD+anti-p75^NTR^ – 0.62 ± 0.03. **(e)** Example tertiary dendrites from CA1 pyramidal neurons of organotypic hippocampal slices immunostained for gephyrin, with or without anti-mBDNF (N-9) treatment before OGD. Scale = 2 µm. **(f)** Quantification of dendritic spines (\*\**p* < 0.001 Two-way ANOVA with Bonferroni multiple comparison test). Total spine density (spines/µm of dendrite): Control – 1.45 ± 0.04 (*n* = 34); Control+anti-mBDNF (N-9) – 1.37 ± 0.03 (*n* = 17); OGD – 0.95 ± 0.05 (*n* = 31); OGD+anti-N-9 – 1.05 ± 0.06 (n>20 cells). **(g)** Quantification of number of gephyrin clusters per confocal stack; consisting of five 512×512 pixel z-planes each (\*\*\**p* < 0.0001, Two-way ANOVA with Bonferroni multiple comparison test; all gephyrin values were normalized to control). Mean number (A.U.): Control – 1.00 ± 0.03 (*n* = 11); Control+N-9 – 0.99 ± 0.04 (*n* = 6 slices); OGD – 0.44 ± 0.04 (*n* = 12); OGD+ anti-mBDNF (N-9) – 0.85 ± 0.06 (*n* = 8). **(h)** Quantification of gephyrin cluster volume (\*\*\**p* < 0.0001, Two-way ANOVA with Bonferroni multiple comparison test). Mean volume (A.U.): Control – 1.00 ± 0.06; Control+N-9 – 1.07 ± 0.06; OGD – 0.66 ± 0.05; OGD+N-9 – 0.88 ± 0.05. Data shown as mean ± s.d.

We next investigated the role of mBDNF in OGD-induced synapse loss. For this we pretreated cultures with anti-mBDNF (N-9, a function blocking antibody) prior to OGD and then quantified the dendritic spines; control sister cultures were processed simultaneously with and without anti-mBDNF treatment. The data revealed a significant downregulation of total dendritic spines in both OGD and anti-mBDNF pretreated OGD slices, compared to control and cultures pretreated with anti-mBDNF (Fig. 3e-f). Similar to our earlier observations, only the mushroom and long-thin subtype of dendritic spines were downregulated 90 min following OGD with or without anti-mBDNF treatment (Fig. 3f). This data suggested to us that mBDNF was not mediating excitatory synapse loss after OGD. In contrast, anti-mBDNF treatment was sufficient to prevent the reduction of gephyrin cluster density in OGD treated slices (Fig. 3g). Reduction of gephyrin cluster volume was also prevented in the presence of anti-mBDNF antibody compared to OGD slices (Fig. 3h). Our results show that mBDNF acts specifically on GABAergic synapses after OGD.

### Blocking ERK1/2 and GSK3β pathways protects gephyrin, but not dendritic spine loss

mBDNF binds with high-affinity to TrkB receptors. It is also known that ERK1/2 and GSK3β pathways are activated downstream of TrkB ^37^. Therefore, we determined whether ERK1/2 and GSK3β signaling cascades downstream of TrkB were activated after OGD to mediate gephyrin cluster reduction at GABAergic terminals. We pretreated slices with pharmacological inhibitors of GSK3β (25 µM GSK3β-IX) and MEK (30 µM PD98059) to prevent activation of these kinases before OGD. Control treated and untreated cultures served for comparative analysis. As expected, inhibiting ERK1/2 and GSK3β pathways did not prevent dendritic spine loss in OGD treated cultures (Fig. 4a, b). Specifically, the mushroom and stubby dendritic spines that were most affected by OGD could not be rescued with ERK1/2 and GSK3β blockade (Fig. 4b). However, at GABAergic synapses gephyrin cluster loss was prevented after OGD in slices pretreated with GSK3β-IX and PD98059, compared to untreated OGD slices (Fig. 4a, c, d).

**Figure 4.**
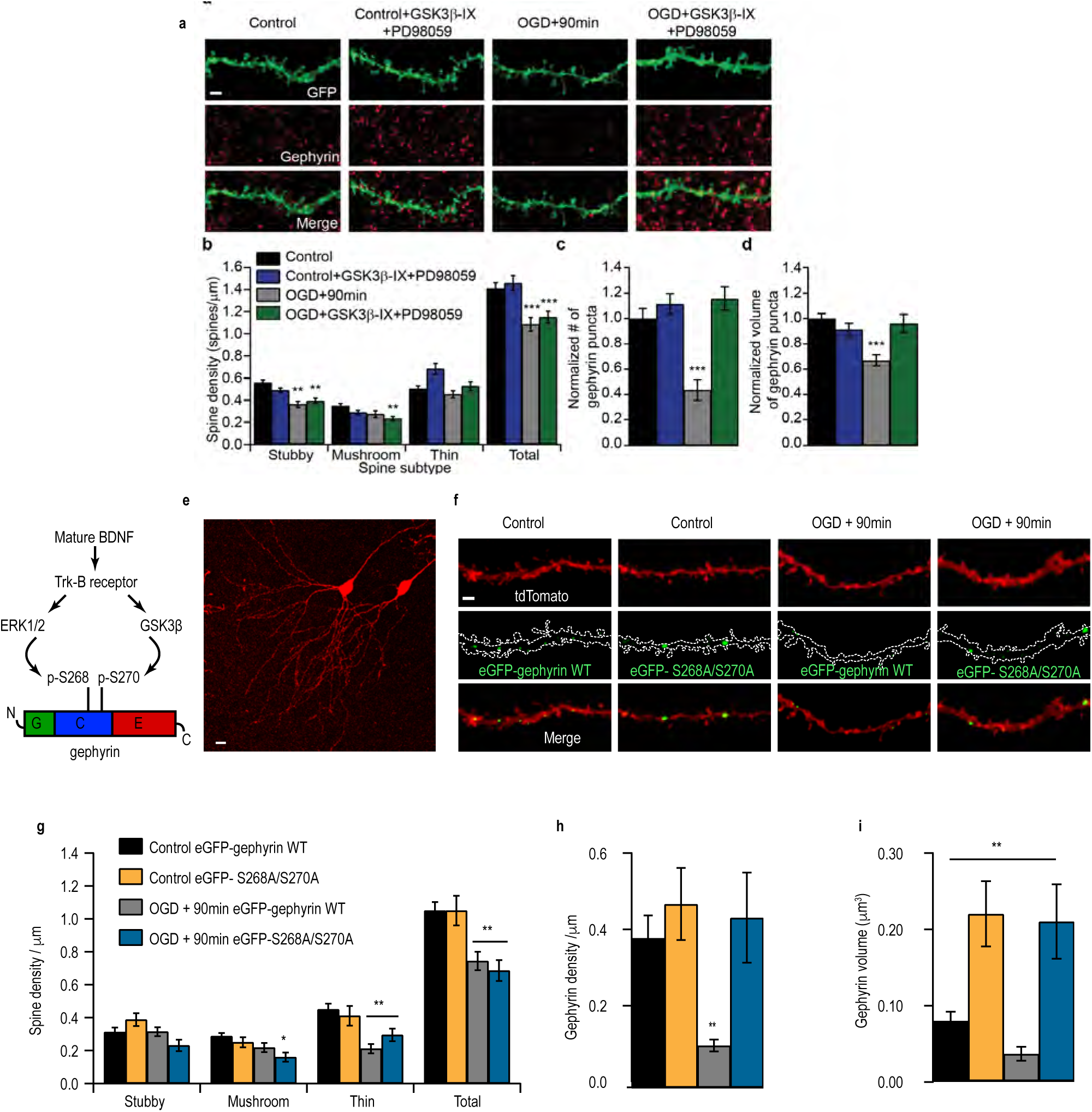
ERK1/2 and GSK3β pathways induce gephyrin degradation but not dendritic spine loss after OGD. **(a)** Example tertiary dendrites from CA1 pyramidal neurons from control and 90 min post-OGD slices, immunostained for gephyrin, with or without pharmacological treatment (GSK3β inhibitor: GSK3β-IX and MEK inhibitor: PD98059). Scale = 2 µm. **(b)** Dendritic spine quantification (\*\**p* < 0.01 Two-way ANOVA with Bonferroni multiple comparison test). Total spine density (spines/µm of dendrite): Control – 1.41 ± 0.06 (*n* = 21); Control+PD+G.-IX – 1.46 ± 0.07 (*n* = 19); OGD – 1.09 ± 0.06 (*n* = 22); OGD+PD+G.-IX – 1.15 ± 0.05 (*n* = 21). **(c)** Quantification of number of gephyrin cluster density (consisting of five 512×512 pixel z-planes each; \*\*\**p* < 0.0001, Two-way ANOVA with Bonferroni multiple comparison test; all gephyrin values were normalized to control). (d) Quantification of number of gephyrin cluster volume. Mean number (A.U.): Control – 1.00 ± 0.08 (*n* = 11); Control+PD+G.-IX – 1.12 ± 0.08 (*n* = 9); OGD – 0.44 ± 0.08 (*n* = 9); OGD+PD+G.-IX – 1.16 ± 0.09 (*n* = 9). **(d)** Quantification of gephyrin puncta volume (\*\**p* < 0.001, Two-way ANOVA with Bonferroni multiple comparison test). Mean volume (A.U.): Control – 1.00 ± 0.04; Control+PD+G.-IX – 0.91 ± 0.05; OGD – 0.67 ± 0.04; OGD+PD+G.-IX – 0.96 ± 0.07. **(e)** Maximum intensity projection of example CA1 pyramidal neurons which had been biolistitally transfected with a plasmid expressing tdTomato. Scale = 10 µm. Neurons were cotransfected with either wildtype or mutant gephyrin. **(f)** Example tertiary dendrites from CA1 pyramidal neurons from control and 90 min post-OGD slices which were transfected with either eGFP-gephyrin_WT_ or eGFP-gephyrin_S268A/S270A_. **(g)** Dendritic spine quantification (\*\**p* < 0.01 Two-way ANOVA with Bonferroni multiple comparison test). Total spine density (spines/µm of dendrite): Control + gephyrin_WT_-GFP – 1.05 ± 0.052 (*n* = 11); Control + eGFP-gephyrin_S268A/S270A_ – 1.05 ± 0.090 (*n* = 12); OGD – eGFP-gephyrin_WT_ 0.74 ± 0.056 (*n* = 14); OGD + eGFP-gephyrin_S268A/S270A_– 0.69 ± 0.064 (*n* = 8); \*\**p* < 0.01, Two-way ANOVA with Bonferroni multiple comparison test). **(h)** Quantification of number of gephyrin cluster density (\*\**p* < 0.01, Two-way ANOVA with Bonferroni multiple comparison test). Density: Control + eGFP-gephyrin_WT_ – 0.38 ± 0.58; Control + eGFP-gephyrin_S268A/S270A_ – 0.47 ± 0.092; OGD + eGFP-gephyrin_WT_ – 0.11 ± 0.015; OGD+ eGFP-gephyrin_S268A/S270A_– 0.44 ± 0.12. **(i)** Quantification of gephyrin cluster volume (\*\**p* < 0.01, Two-way ANOVA with Bonferroni multiple comparison test). Mean volume: Control + eGFP-gephyrin_WT_ – 0. 85 ± 0.012; Control + eGFP-gephyrin_S268A/S270A_ – 0.22 ± 0.043; OGD + eGFP-gephyrin_WT_– 0.046 ± 0.01; OGD+ eGFP-gephyrin_S268A/S270A_ – 0.21 ± 0.049. Data shown as mean ± s.d.

Previously, we have reported that GSK3β phosphorylates gephyrin on serine 270 (ser270) to negatively regulate the number of gephyrin clusters ^38^, and ERK1/2 phosphorylates gephyrin at serine 268 (ser268) to negatively regulate the size of gephyrin clusters ^39^. In order to determine whether phosphorylation of these serine residues were important for OGD-induced gephyrin downregulation, we used biolistic transfection of GFP-tagged gephyrin where serines 268 and 270 were mutated to alanines (gephyrin_S268A/S270A_) into CA1 pyramidal neurons and assessed whether this gephyrin mutant is insensitive to mBDNF-mediated TrkB signaling after OGD. Our analysis showed that gephyrin_S268A/S270A_ mutant is resistant to OGD compared to wildtype gephyrin (gephyrin_WT_) (Fig. 4e, f, h, i). Transgene expression of gephyrin_S268A/S270A_ mutant could not prevent dendritic spine loss after OGD, which is consistent with the data using pharmacological inhibitors that block kinase pathways directly phosphorylating gephyrin at Ser268 and Ser270 respectively (Fig. 4g). Overall, our results identify gephyrin S268 and S270 phosphorylation downstream of TrkB as a determinant for GABAergic synapse loss after OGD.

### The MCAO model *in vivo* recapitulates OGD-induced synapse loss at 24 h post ischemia

In order to confirm our *in vitro* OGD results also occurred in *in vivo* we used MCAO technique, in which an intraluminal filament is used to cause transient ischemia in the fronto-parietal cortex and striatum ^40^, and assayed for glutamatergic and GABAergic synapse loss 24 h post MCAO. MCAO is the most extensively used model in rodents as it produces a reproducible infarct (core and peri infarct area) where pathophysilogical cascades are well described. The peri-infarct area surrounding the core is the site for inflammation, synaptic plasticity and circuit adaptations where structural and functional changes within cortex have been observed in patients after 3 months following an anterior ischemic stroke ^41^. Cell death within the ischemic core renders the tissue fragile for morphology or functional analysis; hence, we restricted our analysis to the penumbra.

We used immunohistochemical staining to assess changes in glutamatergic and GABAergic synapse changes within the peri-infarct area and in the corresponding area contralaterally as comparison. For glutamatergic synapse labeling we chose VGLUT1 to test for glutamatergic presynaptic changes and PSD95 for glutamatergic postsynaptic changes (Fig. 5a, b, c). For GABAergic synapse labeling we chose GAD65/67 to label presynaptic terminals; GABA_A_R γ2 for synaptic receptors and GABA_A_R *α*5 for extrasynaptic receptors (Fig. 5a, d, e, f). Analysis of parietal cortex layer 2/3 (L2/3) 24 h after MCAO did not show any observable loss of PSD95 (Fig. 5c). This result is consistent with previous report that, under ischemic conditions, nNOS interacts with PSD95 to stabilize it at the cell membrane ^42^. Analysis for VGLUT1-positive terminals showed a significant reduction in presynaptic sites in both ipsi- and contra-lateral hemispheres (Fig. 5b).

**Figure 5.**
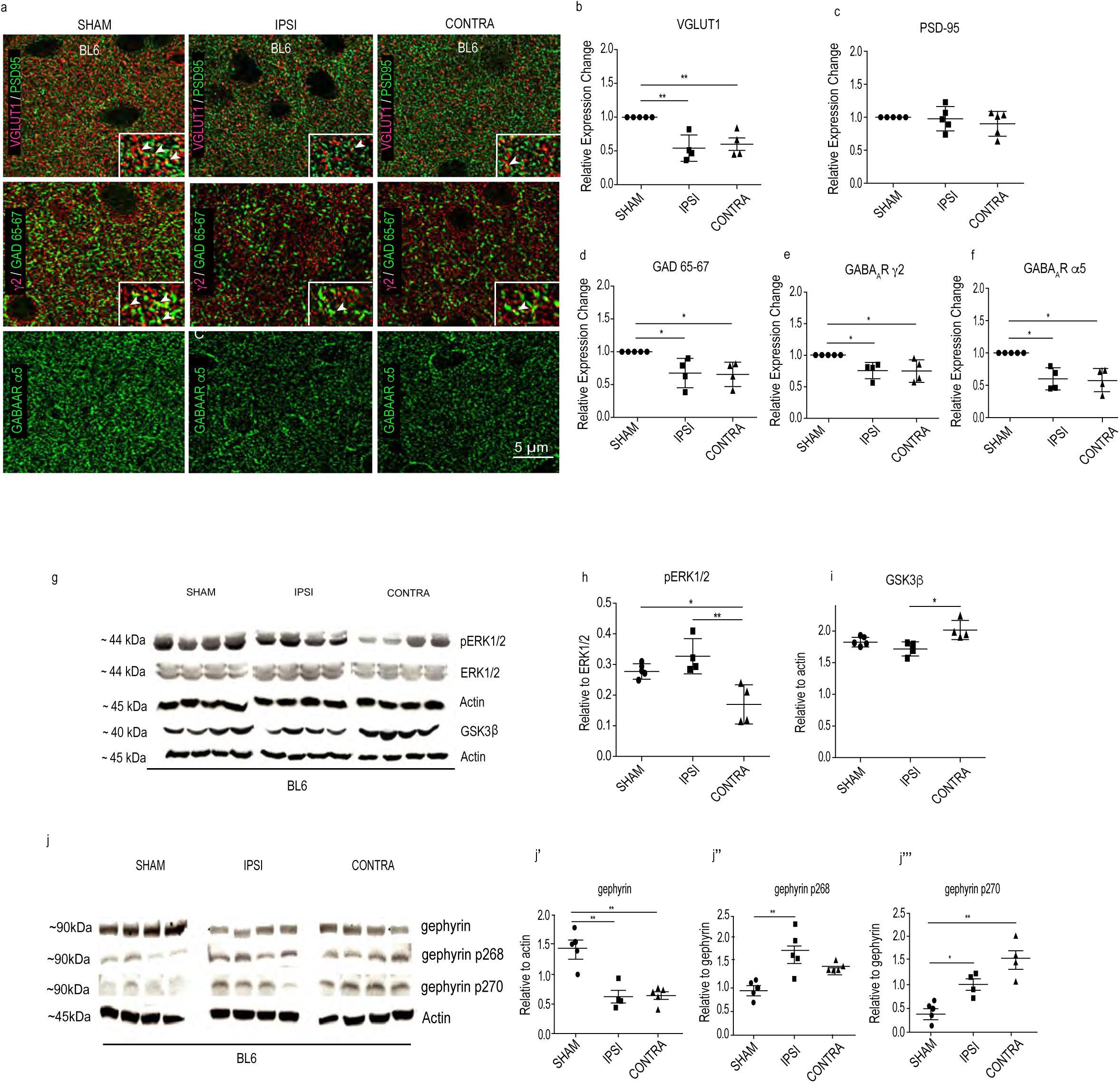
Synapse changes in the peri-infarct area of ipsi-lateral hemispheres 24 h following MCAO. **(a)** Example composite images for glutamatergic synaptic proteins (VGLUT1 and PSD95) and GABAergic synaptic markers (GAD65/67, *γ*2 and *α*5 GABA_A_Rs). **(b)** Quantification for VGLUT1 cluster density (One-way ANOVA, Bonferroni multiple comparison post-hoc test; F(2,49)=11.2; P<0.0001). **(c)** Quantification for PSD95 cluster density (One-way ANOVA, Bonferroni multiple comparison post-hoc test; F(2,49)=0.84; P=0.43). **(d-f)** Quantification for GAD65/67 (One-way ANOVA, Bonferroni multiple comparison post-hoc test; F(2,10)=7.9; P=0.0085), *γ*2 (One-way ANOVA, Bonferroni multiple comparison post-hoc test; F(2,9)=6.4; P=0.018) and *α*5 GABA_A_R cluster density (One-way ANOVA, Bonferroni multiple comparison test; F(2,9)=8.3; P=0.0088). Data shown as mean ± s.d. (n=5 animals); *P<0.05; **P<0.01; ***P<0.001. **(g-i)** WB analysis for phospho-ERK1/2, total ERK1/2, GSK3*β* and actin in sham and 24 h post MCAO samples. **(j-j’’’)** WB analysis for total gephyrin, phospho gephyrin-S268 and phospho gephyrin-S270 levels in BL6 WT sham and 24 h post MCAO mice. Data shown as mean ± s.d. (n=4 animals).

At GABAergic synapses, we saw a significant reduction in GAD65/67 terminals, γ2 subunit-containing synaptic GABA_A_Rs and *α*5 subunit-containing extrasynaptic GABA_A_Rs (Fig. 5d-f). We could not morphologically assess for α1 and α2 subunit-containing GABA_A_Rs in MCAO tissue due to the strong post-fixation protocol that is not condusive for these two antibodies. Therefore, we used Western blot (WB) to examine the expression level of *α*1 and *α*2 subunits in control and MCAO tissue from both ipsi- and contra-lateral hemispheres of parietal cortex L2/3. We found a significant reduction in α1 and α2 GABA_A_R subunit expression after MCAO (Suppl. Fig 2a, b). Overall, our analysis confirms that both ipsi- and contra-lateral hemispheres decrease protein expression of important glutamatergic and GABAergic synaptic markers. Therefore, reduced excitatory and inhibitory synapse markers *in vivo* recapitulate our *in vitro* OGD results showing impaired glutamatergic and GABAergic synaptic transmission.

Blocking effector kinases downstream of TrkB *in vitro* OGD experiments effectively rescued GABAergic synapse loss (Fig. 4c-d) and transgene expression of gephyrin S268A/S270A mutant insensitive to ERK1/2 and GSK3*β* kinases prevented gephyrin cluster loss after OGD (Fig. 4h-i). Hence, we assessed ERK1/2 and GSK3*β* kinase activation levels 24 h after MCAO in fronto-parietal cortex ipsi- and contralaterally in BL6 WT mice. WB analysis for ERK1/2 and its phosphorylated form showed unchanged ERK1/2 levels but significantly increased levels of phosphorylated ERK1/2 in the ipsi- but not the contra-lateral hemisphere at 24 h after MCAO (Fig. 5g, h). WB analysis for GSK3*β* showed increased kinase expression in the contra-lateral hemisphere but not the ipsi-lateral hemisphere at 24 h after MCAO (Fig. 5i). Gephyrin expression level and its phosphorylation at Ser 268 and Ser 270 sites changed at 24 h after MCAO (Fig. 5j- j’’’). WB quantification confirmed that total gephyrin protein levels significantly decrease at 24 h post MCAO (Fig. 5j’). Consistent with our observation of elevated ERK1/2 activation on the ipsi-lateral hemisphere, we observed significantly higher S268 phosphorylation on gephyrin (Fig. 5j’’). Similarly, higher GSK3*β* levels in the contra-lateral hemisphere correlated with significantly higher gephyrin S270 phosphorylation (Fig. 5j’’’). These observations are consistent with our *in vitro* OGD data, and further confirm a role for ERK1/2 and GSK3*β* pathways in directly phosphorylating gephyrin to regulate protein stability after MCAO.

### Synapse loss after MCAO is attenuated in GphnS268A/S270A point mutant mice

To obtain a more direct confirmation for the central role of gephyrin phosphorylation in synapse alterations 24 h post-MCAO, we generated a *Gphn*S268A/S270A global point mutant mouse line using CRISPR/Cas9 (Cyagen, USA). We performed MCAO in *Gphn*S268A/S270A mutant mice and compared synapse plasticity changes in the parietal cortex L2/3 with sham *Gphn*S268A/S270A littermates. We analysed for changes in glutamatergic and GABAergic synaptic markers using immunohistochemical analysis 24 h post-MCAO (Fig. 6). In the *Gphn*S268A/S270A mutant mice, we observed stabilization of excitatory VGLUT1-positive terminals on both ipsi- and contralateral hemispheres 24 h post MCAO (Fig. 6a, b, b’). PSD95 cluster density was also unchanged in the ipsi- and the contra-lateral hemispheres (Fig. 6b’). Analysis of GABAergic synaptic markers showed a significant increase in the GAD65/67 puncta density in both ipsi- and contra-lateral hemispheres (Fig. 6c). Significantly, we did not observe any reduction in γ2- and α5-containing GABA_A_Rs in *Gphn*268/S270A mutant mice 24 h post MCAO (Fig. 6 c’-c’’). Consistent with observed synaptic marker changes, WB analysis for the α1 and α2 subunits in *Gphn*268/S270A mutant mice showed no alterations for these two major GABA_A_R subunits 24 h following MCAO (Suppl. Fig. 2c-d). Taken together, these results substantiate an involvement of ERK1/2 and GSK3*β* as downstream effectors that critically influence gephyrin scaffold stability along with glutamatergic and GABAergic synapse integrity post-MCAO.

**Figure 6.**
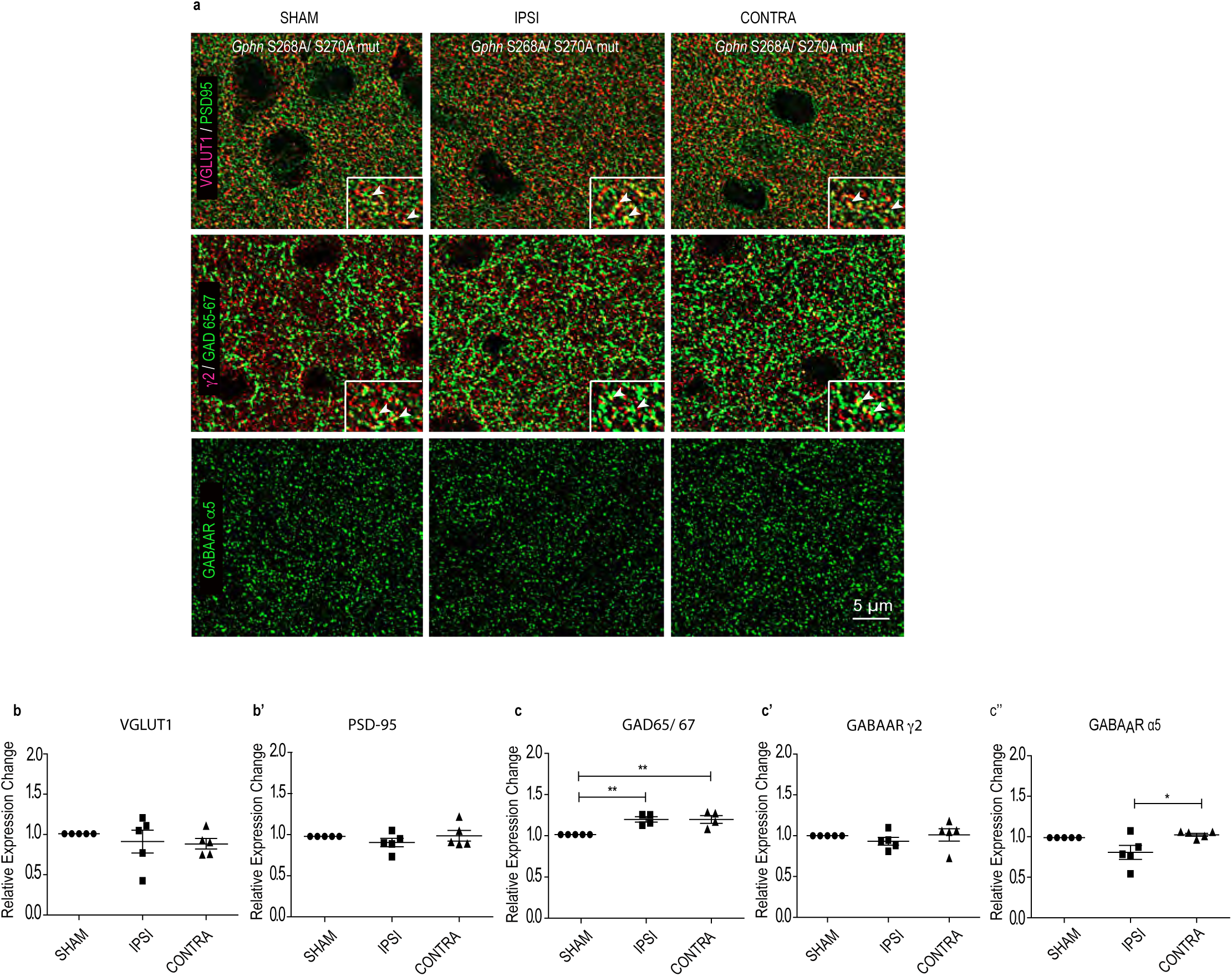
Synapse loss is attenuated in GphnS268A/S270A mutant mice 24 h following MCAO. **(a)** Example images of immunohistochemical stainings from motor cortex L2/3 ipsi- and contra-lateral sides in *Gphn*S268A/S270A mutant mice 24 h following MCAO. Glutamatergic synaptic terminals were visualised using VGLUT1 and PSD95 (see inset). GABAergic synaptic sites were visualised using GAD65/67, *γ*2 and *α*5 GABA_A_Rs. **(b)** Quantification for VGLUT1 (One-way ANOVA, Brown-Forsythe test; F(2,11)=o.35; P=0.70). **(b’)** Quantification of PSD95 cluster density (One-way ANOVA, Brown-Forsythe test; F(2,11)=1.03; P=0.38). **(c)** Quantification for GAD65/67 (One-way ANOVA, Bonferroni multiple comparison test; F(2,9)=11.9; P=0.003). **(c’)** Quantification of *γ*2 (One-way ANOVA, Bonferroni multiple comparison test; F(2,11)=0.60; P=0.56). **(c’’)** Quantification of *α*5 GABA_A_Rs clusters (One-way ANOVA, Bonferroni multiple comparison test; F(2,11)=4.9; P=0.02). Data shown as mean ± s.d. (n=5 animals).

### Microglia induce synapse loss after MCAO

While it is well established that BDNF levels increase after cerebral ischemia, the source of BDNF after stroke remains unclear. Given that *Bdnf* transcripts have been localized within microglia ^43^, we wondered whether microglia contributed to BDNF signaling after stroke. We assessed for BDNF protein changes within ionized calcium-binding adaptor protein-1 (IBA-1) positive cells corresponding to microglia using near super resolution Airy scan microscopy analysis of sham and MCAO BL6 WT mice. Under baseline sham condition, we found low BDNF colocalization within IBA1 positive microglia (Suppl. Fig. 3a, a’). However, following MCAO we could detect elevated BDNF protein within both soma and processes on both ipsi- and contra-lateral hemispheres (Suppl. Fig. 3a). Quantification confirmed an increase in BDNF protein within microglia at 24 h post MCAO (Suppl. Fig. 3a’), implicating elevated BDNF protein translation within microglia in the synaptic pathology of stroke.

To directly examine the role of microglia in BDNF synthesis and synaptic changes after MCAO, we next sought to deplete microglia from the brain using the pharmacological inhibitor PLX5622 that targets colony stimulating factor 1 receptor (CSF-1R) phosphorylation in microglia (Plexxikon Inc. Berkeley, CA 94710). It has been reported that prolonged administration of this drug (1 week) in a formulated chow diet depletes 90% of the microglia from rodent brain ^44, 45^. Replacing the mice on regular diet repopulates microglia cells within 5-7 days. No adverse changes to synapse structure and function, or transcriptional changes have been reported after PLX5622 chow administration ^46^. We confirmed the loss of microglia after PLX5622 chow administration after 7 days (Suppl. Fig. 3b, b’). ^43^

In order to test whether microglia contribute to synaptic changes observed at 24 h post MCAO, we treated BL6 WT with PLX5622 formulated chow or control chow. We then assessed for synapse alterations by staining for synaptic markers and found no significant differences in either glutamatergic synaptic markers (VGLUT1, PSD-95) or GABAergic synaptic markers (GA65/67, α5 and γ2 GABA_A_Rs) between the groups (Suppl. Fig. 3c-h). Similarly, WB analysis to assess α1 or α2 GABA_A_R subunit expression showed no changes at 24 h post-MCAO in mice administered with PLX5622 (Suppl. Fig. 2e, f). In addition, WB analysis to assess total gephyrin or changes in gephyrin phosphorylation at S268 and S270 in PLX5622 treated mice showed no changes (Suppl. Fig. 3i-i’’’). These findings point to the critical role of microglia in mediating synapse loss after MCAO.

### Microglia release BDNF after MCAO to induce synapse loss

BDNF has been shown to play a critical role in the activation of microglia *in vitro*, and increase in BDNF has been tightly linked to pro-inflammation responses ^47, 48^. To specifically evaluate the role of BDNF within microglia in synapse loss following MCAO, we used CX3CR1^ERT2Cre+/-^ mice specifically expressing tamoxifen-inducible Cre recombinase in microglia cells and generated BDNF^flox/flox^ / CX3CR1^CreERT2+/-^ cKO mouse line, thereby preventing *Bdnf* expression selectively in microglia. We used BDNF^wt/wt^ / CX3CR1^CreERT2+/-^ and BDNF^flox/flox^ / CX3CR1^CreERT2+/-^ mice to culture microglia from post-natal day 3 pups and treated with tamoxifen to confirm *Bdnf* gene deletion in microglia cells. qRT-PCR analysis confirmed significant reduction in microglial *Bdnf* mRNA specifically from BDNF^flox/flox^ / CX3CR1^CreERT2+/-^ mice (Suppl. Fig. 4a). Next, we confirmed the loss of BDNF protein from BDNF^flox/flox^ / CX3CR1^CreERT2+/-^ mice *in vivo*. We stained for BDNF and IBA1 in brain slices from BDNF^wt/wt^ / CX3CR1^CreERT2+/-^ and BDNF^flox/flox^ / CX3CR1^CreERT2+/-^ mice. We observed BDNF colocalization in microglia soma and processes in the BDNF^wt/wt^ / CX3CR1^CreERT2+/-^ tissue but not in BDNF^flox/flox^ / CX3CR1^CreERT2+/-^ mice tissue (Suppl. Fig. 4b, b’).

Having confirmed that our cKO mouse line efficiently deletes BDNF in microglia cells, we examined the contribution of microglia *Bdnf* to synaptic changes observed at 24 h post MCAO. For this we assessed changes in synaptic markers (VGLUT1,GAD65/67, GABA_A_R γ2 and GABA_A_R *α*5) in both ipsi- and contra-lateral hemispheres at 24 h post-MCAO compared to BDNF^flox/flox^ / CX3CR1^CreERT2+/-^ sham animals. There was no significant changes between the MCAO and sham groups (Suppl. Fig. 4c-h). We also performed WB analysis to measure α1 and α2 GABA_A_R subunit expression level changes between MCAO and sham group (Suppl. Fig 2g, h). Our analysis confirmed that synaptic receptor expression is unchanged upon *Bdnf* gene depletion from microglia cells post-MCAO. Our earlier observation showed stabilized gephyrin protein levels and no increase in gephyrin phopshorylation at Ser268 and Ser270 after microglia depletion using PLX5622 (Suppl. Fig. 3i-I’’’). In order to confirm if microglial BDNF signaling led to gephyrin protein loss and elevated phosphorylation at Ser268 and Ser270 residues, we performed WB analysis using tissue from BDNF^flox/flox^ / CX3CR1^CreERT2+/-^ sham and MCAO animals (Suppl. Fig. 4i-i’’’). The WB analysis showed that total gephyrin levels and gephyrin phosphorylation at S270 were unchanged, while gephyrin phosporlyation at S268 is reduced post MCAO. Our results uncover a consistent pattern of gephyrin stabilization, reduced gephyrin phosphorylation at Ser268 and Ser270 residues and synapse preservation at 24 h post-MCAO in BL6 mice treated with PLX5622 and BDNF^flox/flox^ / CX3CR1^CreERT2+/-^ transgene mice, suggesting that microglia derived BDNF signals for gephyrin phosphorylation and subsequent synapse downregulation at 24 h post MCAO.

Microglia have been implicated in the rapid engulfment and clearance of synapses following inflammatory brain pathology ^49^.To assess if *Bdnf* gene deletion influenced microglia ramification post MCAO, we stained for IBA-1 in BDNF^wt/wt^ / CX3CR1^CreERT2+/-^ and BDNF^flox/flox^ / CX3CR1^CreERT2+/-^ mice. It has been reported that macrophages infiltrate into the CNS only at day 4 following MCAO ^50^; therefore, IBA-1 positive cells are likely to be resident microglial cells. We performed 3D volume analysis of reconstructed microglia cells from ipsi- and contra-lateral hemispheres (Suppl. Fig. 5a). Quantification showed a significant volume increase in the ipsi- but not the contra-lateral hemisphere in BDNF^wt/wt^ / CX3CR1^CreERT2+/-^ mice, while there was no change in microglia volume in BDNF^flox/flox^ / CX3CR1^CreERT2+/-^ mice 24h post MCAO (Suppl. Fig. 5b). Our data shows significant hypertrophy, indicative of microglial activation, in the ipsi-lateral hemisphere of only BDNF^wt/wt^ / CX3CR1^CreERT2+/-^ mice. The lack of microglial activation in BDNF^flox/flox^ / CX3CR1^CreERT2+/-^ mice suggests that prevention of microglia BDNF release not only preserves synapses, but also prevents alterations in microglia morphology.

### GphnS268A/S270A mutation or Bdnf deletion from microglia reduce brain damage after MCAO

While BDNF has been shown to play the role of pro-survival factor, including microglia activation *in vivo* ^14^, there is evidence to suggest that neuronal activity-dependent exocytosis and/or release from microglia can contibute to sepcific conditions of brain pathology ^51^. To test this, we performed MCAO in BL6 WT, BL6 WT mice treated with PLX5622, *Gphn*S268A/S270A mutant mice or BDNF^flox/flox^ / CX3CR1^CreERT2+^/^-^ cKO (Fig. 7a, b). We used cresyl violet staining to measure the infarct volume across brain sections 24 h following MCAO. Quantification confirmed that in *Gphn*S268A/S270A mutant mice and BDNF^flox/flox^ / CX3CR1^CreERT2+/-^ cKO mice the ischemic tissue damage was significantly reduced as compared to BL6 WT mice. We see a trend in reduced infarct volume in PLX5622 treated mice that does not reach significance. It is likely that ablating microglia completely causes compensatory adaptations in the brain that are not identitical to microglia-specific *Bdnf* gene deletion. These results support our idea that prevention of microglia BDNF release after ischemia or blocking its downstream phosphorylation target gephyrin enhances tissue preservation 24h post-MCAO.

**Figure 7.**
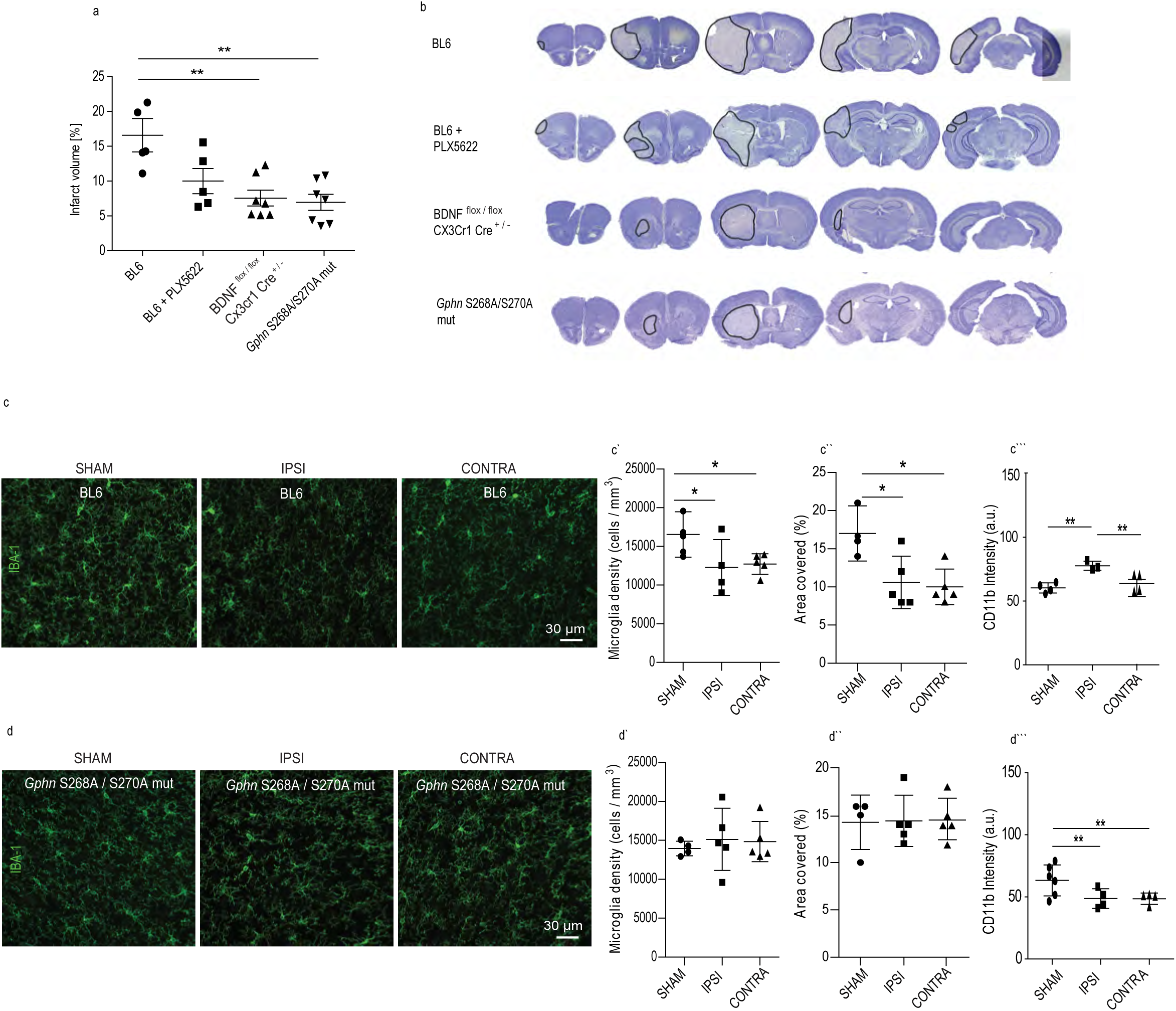
GphnS268A/S270A mutant mice show reduced neuroinflammation 24 h following MCAO. **(a)** Quantification of infarct volume in BL6 WT, BL6 WT treated with PLX5622, BDNF^flox/flox^ / CX3CR1^ERT2Cre+/-^ and *Gphn*S268A/S270A mutant mice at 24 h following MCAO (One-way ANOVA, Brown-Forsythe test; F(3,15)=8.9; P=0.0028). **(b)** Example image of volumetry analysis from 24 h post MCAO tissue sections from BL6 WT, BL6 WT treated with PLX5622, BDNF^flox/flox^ / CX3CR1^ERT2Cre+/-^ and *Gphn*S268A/S270A mutant mice. Results expressed as mean ± s.d. (N=5 or more). **(c)** Example images of BL6 WT mice stained from sham or 24 h post MCAO tissue for microglia markers IBA1 and CD11b (IBA1 shown). **(c’)** Quantification for microglia density in sham animals and ipsi- and contra-lateral sides of MCAO animals. **(c’’)** Quantification microglia area covered using IBAI staining from sham animals, ipsi- and contra-lateral sides of MCAO animals. **(c’’’)** Quantification for CD11b intensity in sham animals, ipsi- and contra-lateral sides after MCAO animals. **(d)** Example images of *Gphn*S268A/S270A mutant mice stained from sham or 24 h post MCAO tissue for microglia markers IBA1 and CD11b (IBA1 shown). **(d’)** Quantification for microglia density in sham animals and ipsi- and contra-lateral sides of MCAO animals. **(d’’)** Quantification microglia area covered using IBAI staining from sham animals, ipsi- and contra-lateral sides of MCAO animals. **(d’’’)** Quantification for CD11b intensity in sham animals, ipsi- and contra-lateral sides after MCAO animals. Results expressed as mean ± s.d. (n=5 animals); *P<0.05 (One-way ANOVA, Bonferroni multiple comparison test).

*In vivo* inflammatory processes play a key role in tissue damage and repair. In response to inflammation, microglia acquire properties for reactive species generation and inflammatory cytokine production, and are therefore thought to be principal drivers of pro-inflammatory responses ^52, 53^.We have demonstrated above, that depletion of microglia-derived BDNF release reduces MCAO-induced increase in microglia volume (Suppl. Fig. 5). Given that *Gphn*S268A/S270A mutant mice show enhanced tissue preservation 24h post-MCAO, we anticipated gephyrin scaffold stabilization to reduce microglia activation after MCAO. To assess microglia properties, we first examined the density of resident microglia at baseline and 24 h post MCAO in the brains of BL6 WT and *Gphn*S268A/S270A mutant mice (Fig. 7c, d). Quantification of IBA-1 positive microglia showed a significant reduction of microglia density from both ipsi- and contra-lateral hemispheres after MCAO in BL6 WT mice (Fig. 7c’). Similarly, analysing for area covered by the microglia cells showed significant reduction in both ipsi- and contra-lateral sides (Fig. 7c’’). These observed changes are consistent with the report showing microglia migration changes after stroke ^53^. To confirm inflamation and microglia activation status at 24 h post-MCAO we stained for the activation-state marker CD11b in microglial cells. Quantification for CD11b intensity showed elevated levels only on the ipsi- but not the contralateral side in BL6 WT mice (Fig. 7c’’’). Analysis of microglia density in *Gphn*S268A/S270A mutant mice showed no changes between sham and MCAO samples (Fig. 7d’). Similarly, there was no change in the area covered (Fig. 7d’’). If indeed gephyrin scaffold stabilization leads to reduced microglia activation, we anticipated less activation of microglia in *Gphn*S268A/S270A mutant mice after MCAO. Analysis for CD11b intensity was not elevated after MCAO in the *Gphn*S268A/S270A mutant mice (Fig. 7d’’’). Taken together, our data shows that in *Gphn*268/S270A mutant mice, the mutation selectively blocks MCAO-induced microglia activation and reduces ischemic tissue damage.

### Bdnf deletion from microglia or GphnS268A/S270A mutation prevent BDNF increase

If gephyrin scaffold stability contributes towards microglia activation after MCAO, then it should be reflected in proBDNF and mBDNF level changes in our different mouse lines. As a first step, we assessed for differences in the baseline level of proBDNF and mBDNF across our different mouse lines. WB analysis showed that BL6 WT mice treated with PLX5622 had elevated levels of proBDNF and BDNF^flox/flox^ / CX3CR1^CreERT2+/-^ mice exhibited significantly lower levels of proBDNF, while others had levels similar to BL6 WT control (Suppl. Fig. 6a, a’). The analysis of mBDNF across different mice lines showed no significant differences (Suppl. Fig. 6a, a’’).

Once we established the baseline differences in proBDNF and mBDNF expression across mice lines used in our study, we went on to compare intra-mouse changes in proBDNF and mBDNF within ipsi- and contra-lateral hemispheres 24 h post-MCAO. In BL6 WT mice, we observed a significant increase in both pro- and mature- forms of BDNF specifically in the ipsi- lateral side; there was a small increase in the contra-lateral side but it was not statistically significant (Suppl. Fig. 6b-b’’). In BL6 WT mice treated with PLX5622, we saw a significant reduction of pro-BDNF levels in both hemispheres 24h post MCAO compared to sham group (Suppl. Fig. 6c-c’). This reduction suggests that resident microglia themselves are an important source of proBDNF or alternatively that microglia signal for proBDNF release elsewhere following MCAO. Levels of mBDNF showed no changes within both ipsi- and contra-lateral hemispheres in PLX5622 treated mice (Suppl. Fig. 6c’’). In BDNF^flox/flox^ / CX3CR1^CreERT2+/-^ cKO, although proBDNF levels were significantly lower than BL6 WT animals at baseline, there was no significant difference within the genotype after MCAO (Suppl. Fig. 6d, d’). However, mBDNF that was at similar levels to BL6 WT, showed a significant reduction in both ipsi- and contra-lateral hemispheres after MCAO within the genotype (Suppl. Fig. 6d-d’’). We then assessed BDNF level changes in the *Gphn*S268A/S270A mutant mice. Quantification of the WB showed that gephyrin scaffold stabilization prevented the ipsilateral increase in proBDNF and mBDNF levels observed in BL6 WT animals after MCAO (Suppl. Fig. 6e-e’’). These observations suggests that 1) microglia are an important source of BDNF after MCAO and that 2) microglial proBDNF and mBDNF release is influenced by the phosphorlyation status of gephyrin at S268 and S270 following MCAO.

We looked for direct evidence linking gephyrin phosphorylation at Ser268 and Ser270 contributing to BDNF changes within microglia after MCAO. In this regard, we stained for BDNF and IBA-1 and used near super resolutin Airy scan microscopy to quantify BDNF protein changes within microglia of BDNF^wt/wt^ / CX3CR1^CreERT2+/-^, BDNF^flox/flox^ / CX3CR1^CreERT2+/-^ and *Gphn*S268A/S270A mutant mice (Fig. 8a-c’). Quantification of BDNF intensity within IBA-1 cells confirmed increase within ipsi- and contra-lateral hemispheres of BDNF^wt/wt^ / CX3CR1^CreERT2+/-^ mice (Fig. 8a’). However, there was no change in BDNF intensity within microglia of BDNF^flox/flox^ / CX3CR1^CreERT2+/-^ mice or *Gphn*S268A/S270A mutant mice (Fig. 8b’, c’). These results indeed confirm that gephyrin phosphorylation regulates BDNF levels within microglia to initiate release and downstream signaling that ultimately lead to synapse loss following cerebral ischemia (see model Fig. 8d).

**Figure 8.**
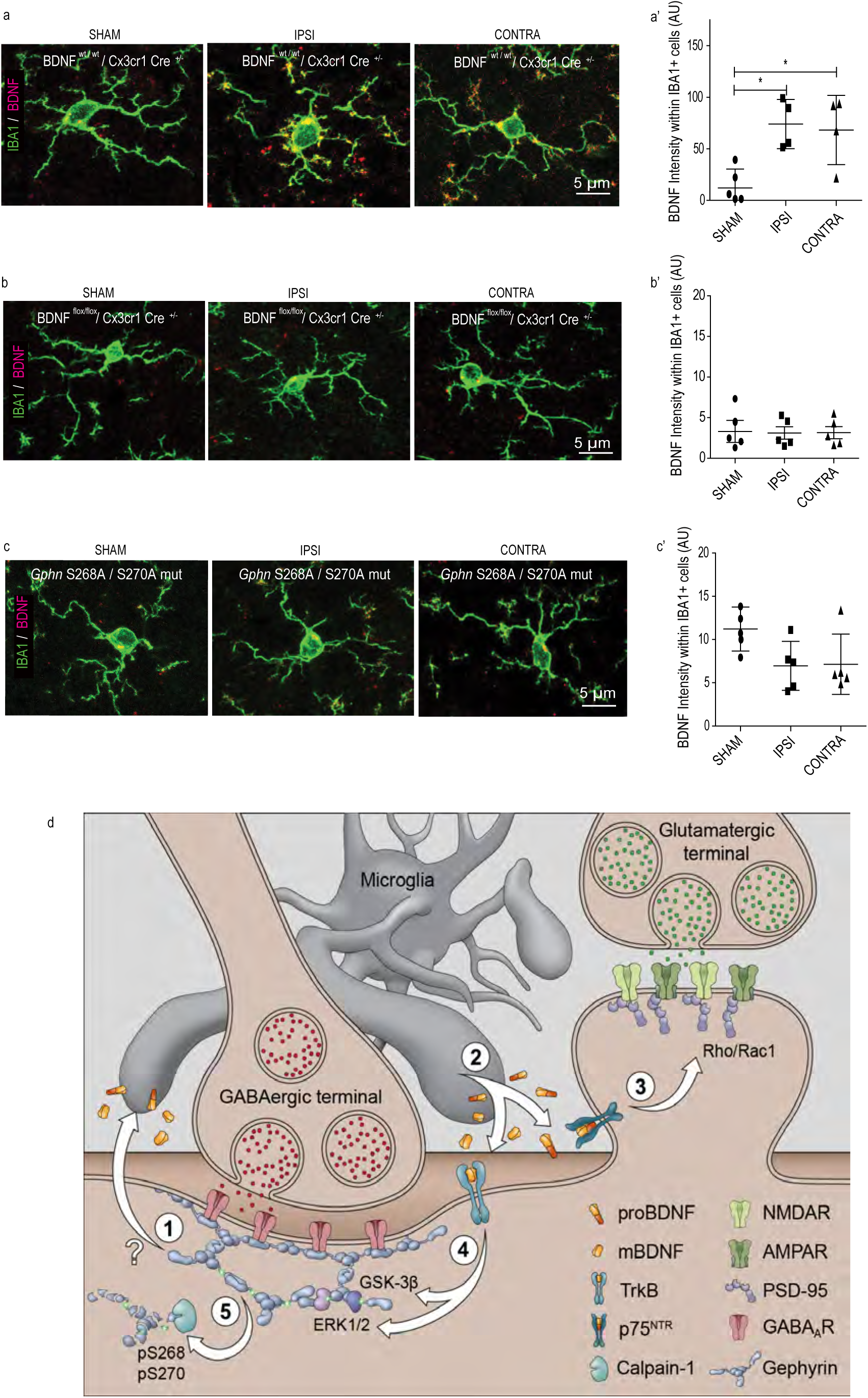
GphnS268A/S270A mutation affects BDNF levels in microglia after MCAO. **(a)** Example images from BDNF^wt/wt^ / CX3CR1^ERT2Cre+/-^ mice stained from sham or 24 h post MCAO tissue for microglia markers IBA1 and BDNF (composite shown). **(a’)** Quantification for BDNF intensity within IBA1 positive cells in BDNF^wt/wt^ / CX3CR1^ERT2Cre+/-^ sham animals and ipsi- and contra-lateral sides of MCAO animals. **(b)** Example images from BDNF^flox/flox^ / CX3CR1^ERT2Cre+/-^ mice stained from sham or 24 h post MCAO tissue for microglia markers IBA1 and BDNF. **(b’)** Quantification for BDNF intensity within IBA1 positive cells in BDNF^flox/flox^ / CX3CR1^ERT2Cre+/-^ sham animals and ipsi- and contra-lateral sides of MCAO animals. **(c)** Example images from *Gphn*S268A/S270A mutant sham animals and ipsi- and contra-lateral sides of MCAO animals. **(c’)** Quantification for BDNF intensity within IBA1 positive cells in *Gphn*S268A/S270A mutant sham animals and ipsi- and contra-lateral sides of MCAO animals. **(d)** Overview of pathway(s) regulating synaptic downregulation downstream of microglia release proBDNF and mBDNF; **(1)** an unknown signal from GABAergic postsynaptic sites facilitates microglia hyper reactivity after MCAO; **(2)** activated microglia produce and release proBDNF and mBDNF to act on their cognate receptors p75^NTR^ and TrkB respectively; **(3)** proBDNF via p75^NTR^ activates RhoA/Rac1 pathway for actin remodeling and spine loss, triggering microglia aided stripping of VGLUT terminals; **(4)** ERK1/2 and GSK3*β* pathways are activated downstream of TrkB receptors to phosphorylate gephyrin at S268 and S270 residues respectively; **(5)** gephyrin phosphorylation at S268 and S270 leads to gephyrin scaffold loss via calpain-1 clevage to facilitate GABA_A_R internalization and microglia aided displacement of GABAergic terminals. This mechanism outlines how excitatory and inhibitory synapses are lost at both ipsi- and contra-lateral sides 24 h post ischemia.

## Discussion

This study reveals a direct connection between microglia, BDNF signaling and gephyrin phosphorylation as a key pathway regulating tissue integrity and synapse loss after ischemic stroke. Preventing BDNF release from microglia or preventing gephyrin phosphorylation at Ser 268 and Ser 270 are equally protective against synapse loss within 24 h post ischemic injury. Specifically, we demonstrate that (a) microglia released proBDNF and mBDNF act through p75^NTR^ and TrkB receptors respectively to facilitate glutamatergic and GABAergic synapse loss after ischemia; (b) ERK1/2 and GSK3*β* kinase phosphorylate gephyrin at Ser 268 and Ser 270 downstream of TrkB for GABAergic synapse downregulation; (c) *in vivo* microglia-specific *Bdnf* gene deletion or expression of gephyrin phospho-null S268A/S270A mutant protect against synapse loss at 24 h post-MCAO; (d) *Gphn*S268A/S270A mutant mice prevent microglia activation and synapse loss at 24 h post-MCAO; (e) the mouse lines tested in our study consistently indicate similar changes within both ipsi- and contra-lateral hemispheres which is rarely addressed in stroke studies. Taken together, microglia activation and BDNF secretion are tightly coupled to glutamatergic and GABAergic synapse integrity in our model of ischemia-induced brain injury.

### GABAergic system in ischemia

Our results concur with previous observations in the gerbil where following transient cerebral ischemia (24 h) GABA_A_Rs are downregulated ^11^. In mice, it was reported that tonic GABA currents are increased in the peri-infarct area 3 days after ischemia. An impairment in GABA transporter GAT3/GAT4 function was shown to contribute towards upregulation of extrasynaptic *α*5 and *δ* GABA_A_R tonic current ^54^. In this study, we report a reduction in synaptic GABA_A_R subunits *α*1, *α*2, *γ*2 and also extrasynaptic *α*5 subunit expression at 24 h after ischemia (Fig. 5; Suppl. Fig. 2). In addition, we report that presynaptic GAD65/67 terminals are also significantly reduced at 24 h after ischemia. Importantly, our data indicates that by stabilizing GABAergic synaptic and extrasynaptic transmission, one can reduce ischemic brain damage (Fig. 7a). It has also been reported that inverse agonists specific for α5-subunit-containing extrasynaptic GABA_A_ receptors administered 4 days after stroke promotes early stroke recover ^54^. Hence, it is conceivable that increase in extrasynaptic GABA_A_Rs observed at day 4 post ischemia is a homeostatic response to the early reduction in GABA and synaptic GABA_A_Rs. Stabilizing the gephyrin scaffold could thus explain the neuroprotection observed in *Gphn*268A/S270A mutant mice at 24 h post ischemia.

### Postsynaptic scaffold stability after ischemia

In WT mice, 24 h post MCAO gephyrin protein levels are reduced (Suppl. Fig. 3e); however, we observe no change in PSD95 clusters, as a proxy for glutamatergic post-synaptic sites (Fig. 5c). Previous studies have revealed that after ischemia nNOS interaction with PSD95 stabilizes the protein at postsynaptic sites ^42^. Interestingly, gephyrin has also been reported to interact with nNOS ^55^. However, it remains to be tested whether interaction with nNOS is influenced by ERK1/2 and GSK3*β* phosphorylation of gephyrin. It is possible that reduced gephyrin expression after MCAO might make more nNOS available for PSD95 interaction and stabilization. If neuroinflammation is the trigger for nNOS activation, our data shows reduced inflammation in *Gphn*S268A/S270A mutant mice and BDNF^flox/flox^; CX3CR1^CreERT2+/-^ cKO mice. Therefore, BDNF signaling could increase neuroinflammation by activating nNOS after ischemia.

### proBDNF and mBDNF signaling regulate synapse downregulation

At the neuromuscular junction, proBDNF and mBDNF elicit opposite effects by promoting axon retraction through activation of p75^NTR^ on presynaptic site or potentiate synapse through TrkB activation at the postsynaptic site respectively. High-frequency neuronal activity controls the ratio of extracellular proBDNF/mBDNF by regulating the secretion of extracellular proteases, serving as a reward signal to stabilize synaptic contacts and strengthen neurotransmission ^56^ However, within hippocampal neurons, proBDNF has been reported to activate p75^NTR^ localized in dendritic spines of CA1 neurons and enhance NR2B-dependent LTD ^57^.

In the current study, we present evidence for proBDNF and mBDNF in glutamatergic and GABAergic synapse down regulation after ischemia. p75^NTR^ lack intrinsic enzymatic activity and activate signal transduction pathway by associating with adaptor proteins that are distinct from TrkB signaling cascade ^58^. Consistent with this literature, our data shows the specificity for proBDNF and p75^NTR^ signaling for spine downregulation and mBDNF and TrkB for gephyrin regulation at GABAergic postsynaptic sites (Fig. 3). Our study has important implications. First, we uncover a synaptic plasticity function for proBDNF and p75^NTR^ in the ischemic brain, which is in marked contrast to its role in regulating neuronal apoptosis. Second, our results show that TrkB and downstream pathways (ERK1/2 and GSK3*β*) specifically influence the stability of shaft synapse and of not spine synapses (Fig. 4).

### Microglia as a source of BDNF in ischemia

Deletion of BDNF from specific subpopulations of neurons has revealed that both presynaptic and postsynaptic BDNF contributes to specific aspects of LTP. For example, presynaptic BDNF was documented to increase the strength of LTP, while postsynaptic BDNF facilitates LTP maintenance ^59^. In addition, BDNF release from dendritic spines can activate NMDA and TrkB receptors within the same release site to influence structural plasticity ^60^. At GABAergic terminals, time duration of exogenous BDNF exposure has opposite effects on GABA_A_R and gephyrin clustering. In hippocampal neuron culture, short-term (5 min) BDNF application inhibits GABA_A_R internalization through phosphoinositide-3 kinase (PI-3 kinase) and PKC pathways (Jovanovic et al., 2004). However, long-term (90 min) BDNF application reduces GABA_A_R and gephyrin clustering ^20^. Presynaptically, BDNF regulates GAD65 mRNA expression through the recruitment of ERK pathway, leading to cAMP-response element (CRE)-binding protein (CREB) activation ^61^.

To add to this complexity, BDNF is not only found in neurons but also expressed in both astrocytes and microglia ^43, 62^. Within the spinal cord circuit, BDNF activates TrkB in lamina I neurons to downregulate KCC2 (Chloride potassium symporter), thereby increasing intracellular chloride concentration and reversing GABAergic inhibition to cause neuronal depolarization ^48^. The resulting hyperexcitability of neurons contributes to mechanical hypersensitivity. Microglia-specific *Bdnf* knockout reduces PNI-induced pain ^63^ In the current study, we report microglia as the major source of BDNF after ischemia for glutamatergic and GABAergic synapse regulation. We observed elevated proBDNF in PLX5622 treated mice in sham condition, which suggests that microglia ablation at baseline could trigger proBDNF synthesis from other cell-types as compensatory mechanism (Suppl. Fig. 6 a’). Interestingly, proBDNF levels reduce significantly in PLX5622 treated animals after MCAO, indicating that microglia play a pivotal role in *de novo* BDNF synthesis, we also do not know how different metalloproteases are affected after microglia depletion. The release of BDNF from microglia has been linked to purinergic receptor P2X4R activation, causing disinhibition of pain-transmitting spinal lamina I neurons ^64, 65^. The expression of P2X4R increases after exposure to proinflammatory cytokines such as INF-*γ* ^66^. Microglia can sense extracellular ATP through P2Y12R to migrate ^67^, or P2X7R to be activated ^68^. Hence, it is possible that within 24 h post ischemia inflammation activates P2X4R expression to promote BDNF protein synthesis and release.

### Gephyrin phosphorylation influences microglia activation

During acute ischemic stroke natural killer (NK) cells infiltrate periinfarct areas of the brain to promote inflammation (e.g. microglia activation) and neuronal damage. Interestingly, depletion of NK cells within the first 12 h after MCAO attenuates neurological deficits and infarct volume ^69^. Here, we report that microglia depletion using PLX5622 (Suppl. Fig 5) and *Bdnf* gene deletion specifically from microglia (Suppl. Fig. 4) prevent glutamatergic and GABAergic synapse loss and reduce the extent of infarct volume (Fig. 7a) after ischemia. Hence, it is possible that NK cells influence infarct development through microglia activation and local BDNF release to trigger glutamatergic and GABAergic synapse loss. Furthermore, in addition to activation of microglia, we uncover activation of ERK1/2 and GSK3*β* pathways (Fig. 5g-i) 24 h post ischemia. As a direct target of ERK1/2 ^39^and GSK3*β* ^38^, gephyrin phosphorylation at Ser268 and Ser270 sites are significantly increased after ischemia, while total gephyrin level is decreased (Fig. 5 j-j’’’). Stabilization of gephyrin clusters through the expression of S268A/S270A mutant in hippocampal slice culture selectively prevents GABAergic synapse loss after ischemia (Fig. 4h, i). In *Gphn*S268A/S270A mutant mice both glutamatergic and GABAergic synapse loss can be prevented (Fig. 6). This suggests to us that mechanisms activated after ischemia *in vivo* might be similar to mechanisms operational in organotypic slice cultures after OGD.

We reveal a link between gephyrin phosphorylation at Ser268 and Ser270, microglia activation and BDNF secretion (see model; Fig. 8d). Within this model, a currently unknown signal originating at GABAergic postsynaptic sites would activate microglia to synthesize and release proBDNF and mBDNF for downstream signaling after MCAO. This is based on our data showing BDNF protein levels do not change after MCAO in *Gphn*268A/S270A mutant mice, which suggests that gephyrin scaffold stability controls microglia activation. The glutamatergic spine synapse collapse through p75^NTR^ signaling and GABAergic synapse downregulation via gephyrin S268 and S270 phosphorylation could facilitate microglia aided stripping of VGLUT and VGAT terminals. It has been reported that microglia physically displace GABAergic presynaptic terminals after lipopolysaccharide induced inflammation ^70^. Together, our data identifies the mechanistic basis for silencing neurotransmission at the initial 24 h post-MCAO.

## Experimental Procedures

### Ethics Statement

All animal handling procedures were carried out consistent with guidelines set by the Canadian Council on Animal Care, the European Community Council Directives of November 24, 1986 (86/609/EEC) and approved by the cantonal veterinary office of Zurich (ZH219/15). All procedures were approved by the Animal Resource Committee of the School of Medicine at McGill University and are outlined in McGill University Animal Handling Protocol #5057.

### Hippocampal Slice Cultures and Oxygen-Glucose Deprivation (OGD)

We have chosen to study the hippocampus as it possesses a unique unidirectional network that is preserved within organotypic cultures ^71^, making it an ideal candidate to study microcircuitry remodeling. Organotypic hippocampal slices were prepared using the roller-tube method, as previously described ^71^ with transgenic mice expressing membrane-targeted MARCKS-enhanced GFP under the Thy-1 promoter in a subpopulation of CA1 cells ^72^. We used an established model of OGD as a model of ischemia ^73^, briefly, mature organotypic hippocampal slices were placed in a glass dish containing glucose-free Tyrode’s solution (in mM: NaCl, 137; KCl, 2.7; CaCl_2_, 2.5; MgCl_2_, 2; NaHCO_3_, 11.6; NaH_2_PO_4_, 0.4; pH 7.4) containing: 2 mM 2-Deoxyglucose, 3 mM sodium azide and 8 mM sucrose, for 4-5 min, and were then returned to culture media for 90 min, 24 h or 1 week. Control slices were exposed to Tyrode’s solution (in mM: NaCl, 137; KCl, 2.7; CaCl_2_, 2.5; MgCl_2_, 2; NaHCO_3_, 11.6; NaH_2_PO_4_, 0.4; glucose, 5.6; pH 7.4) for 4-5 min and returned to culture media.

### Mouse lines

All procedures fulfilled the ARRIVE guidelines on experimental design, animal allocation to different experimental groups, blinding of samples to data analysis and reporting animal experiments. Littermates from heterozygous breedings were used within similar genotypes and inbred animal backgrounds. We compared results within same genotypes. The *Gphn*S268A/S270A mutant mouse was generated using CRISPR/cas9 (Cyagen, USA) in BL6 background. B6.129P2(C)-Cx3cr1tm2.1(cre/ERT2) (Stock 020940) ^74^mice and Bdnftm3Jae or BDNF^Tg^ (Stock 004339) ^75^ were obtained from Jackson Laboratory to generate BDNF^wt/wt^ / CX3CR1^ERT2Cre+/-^ and BDNF^flox/flox^ / CX3CR1^ERT2Cre+/-^cKO lines. The mice were injected (i.p) on four consequtive days with tamoxifen dissolved in corn oil (Sigma H-6278; 1mg/ day) to induce Cre expression at 4 weeks followed by sham or MCAO surgery at 8-9 weeks of age. Animals were random assigned and both genders were used for both conditions. The PLX5622 treatment for microglia depletion followed recommended company dose (1200mg of active form of PLX5622/kg of chow).

### Middle cerebral artery occlusion (MCOA) model

Wild type C57Bl6/J-Crl1 mice were purchased from Charles River Laboratories (Germany), the *Gphn*S268A/S270A mutant, BDNF^Tg/Tg^ / CX3CR1^ERT2Cre+/-^ were bred in house. At 4 weeks age animals were randomly allocated to groups. The transient occlusion of the middle cerebral artery was conducted at 8-9 weeks age using the filament model as described previously ^76^. Briefly, anaesthesia was induced using 3% isoflurane in an oxygen/air (1:4) mixture and maintained at 2% Isoflurane. The area around the neck was shaved, disinfected and an incision was made along the midline. The common and the external carotid artery were isolated and ligated. A silicon rubber filament (Doccol, USA, Lot 701956RE) was inserted into the internal carotid artery to block the middle cerebral artery. The filament remained in place for 30min before reperfusion was allowed by withdrawing it. During occlusion the mouse was placed in a preheated (30°C) recovery box and allowed to recover from anaesthesia. After reperfusion, the internal carotid artery was ligated to prevent bleeding and the wound was sutured. Sham operation involved identical surgical procedures, but the filament was immediately withdrawn after insertion. A total volume of 500µl of Buprenorphine (0.03mg/g) with Saline was injected after surgery and consecutive after 4h and 8h s.c. The mouse was kept for 2h in the recovery box and then placed back into its home cage. Mashed food and food pellets were placed on the cage bottom to encourage food uptake. The described lesion paradigm caused an extensive damage to the unilateral basal ganglia and the adjacent neocortex. For clinical scoring, mice were examined for forelimb flexion and body posture maintenance using the Bederson score [as described in https://www.ahajournals.org/doi/pdf/10.1161/01.STR.17.3.472]. Animals were excluded from the studies when they fulfilled one of the following criteria: prolonged surgery time (>15 min); no reflow after filament withdrawal; Clinical scoring of 0; seizures during/after MCAO or dead before experimental endpoint. A total of 7 animals were exluded (3 due to clinical scoring of 0 following MCAO and 4 due to seizures during/after MCAO).

### Cresyl violet staining

Cresyl violet staining was performed 24h after of reperfusion and infarct volume was assessed as percentage of the affected hemisphere. Five 20µm thick coronal sections taken at Bregma +2.8, +1.54, +0.14, −1.94 and −4.6mm, were stained with cresyl violet using vendor protocol and later digitalised using a Zeiss Axio Scan.Z1 at 5x magnification and lesions were determined using Zen Software (Zeiss). The person analysing was blinded to the treatment groups. Cerebral lesion volume was calculated summing up the volume of each section, while corrected for oedema (group numbers >4).

### Immunofluorescence and Confocal Microscopy

#### Hippocampal organotypic slices

90 min after OGD induction, hippocampal slices were fixed for 1 h at room temperature in 4% paraformaldehyde in 0.1 M phosphate buffer (PB), pH 7.4. Slices were then washed 5× in 0.1 M PB, permeabilized in 0.4% Triton X-100 and blocked with 1.5% heat inactivated horse serum overnight at 4°C. 1:500 anti-Gephyrin (Synaptic Systems) was incubated for 5 days at 4°C in permeabilizing buffer and washed 5× with 0.1 M PB containing 1.5% heat inactivated horse serum. Slices were incubated for 3 h at room temperature with 1:250 anti-mouse DyLight 650 (Jackson ImmunoResearch, Burlington, ON, Canada) secondary antibody diluted in 0.1 M PB containing 1.5% heat inactivated horse serum. Following 5× washes with 0.1 M PB containing 1.5% heat-inactivated horse serum, slices were mounted with fluorescent mounting medium (DAKO, Mississauga, ON, Canada) onto microscope slides.

Slices were imaged using a Leica TCS SP2 scan head (Leica Microsystems) on a Leica DM6000 B upright microscope, equipped with HCX PL APO 63× NA 1.4 oil immersion objective using a 543 nm HeNe laser line. Image stacks were collected at Z = 0.3 µm and averaged 2-3 times to improve signal-to-noise ratio. For quantification, image stacks were obtained with identical parameters (laser intensity, filters, pinhole size, photomultiplier tube gain and offset). Representative images are maximum intensity projections of 5 sections from each stack.

### Immunohistochemical imaging of brain slices

24 h following MCAO, P60-70 (male or female) mice were anesthetized by intraperitoneal pentobarbital injection (Nembutal; 50 mg/kg) and perfused transcardially with ice-cold oxygenated ACSF ^77^, pH 7.4, for 2min. Brains were immediately fixed in 4% PFA for 3 h at 4°C. After rinsing in PBS, brains were incubated in 30% sucrose (in PBS) at 4°C over-night. 45µm- thick coronal sections were cut from frozen blocks using a sliding microtome (HM400; Microm) and stored at -20°C in antifreeze solution. After 3 times 10 min washes in Tris-Triton Solution (50mM Tris, 150mM NaCl, 0.05% Trito X-100, pH 7.4), sections were incubated in primary antibody solution (50mM Tris, 150mM NaCl, 0.4% Triton X-100, 2% normal goat serum, pH 7.4) at 4°C overnight. Primary antibodies are listed in Table 1. Sections were washed 3 x 10min in Tris-Triton Solution and incubated in secondary antibody solution (50mM Tris, 150mM NaCl, 0.05% Triton X-100, 2% normal goat serum, pH 7.4) for 1 hour at room temperature with secondary antibodies raised in goat. All secondary antibodies used were diluted 1:500. Antibodies conjugated to AlexaFluor-488 and AlexaFluor-647 were purchased from Invitrogen, while antibodies conjugated to Cy3 were purchased from Jackson ImmunoResearch Laboratories. Sections were washed 3 times 10 min in Tris-Triton Solution and mounted on gelatin-coated slides using Fluorescence Mounting Medium (Dako). Z-stack images (4 optical sections, 0.75µm step size) were recorded of all sections using confocal laser scanning microscopy (LSM 700, Carl Zeiss). Images were taken using a 40x objective with a numerical aperture of 1.4, and pixel size of 112 nm^2^. Three juxtaposed images of the motor cortex layer 2/3 were taken on the ipsi- and contra- lateral hemispheres. To reduce variability, multiple images were captured from 3 sections per mouse and total of 4-5 mice were analyzed per condition/genotype. Cluster density values were averaged from these sections. All imaging parameters were kept constant between MCAO and sham animals. For cluster analysis, a custom Python-script using the ImageJ image-processing framework was used. The script can be used as a plugin and is openly available on a github repository (https://github.com/dcolam/Cluster-Analysis-Plugin). Representative example images were processed using ImageJ. Statistical tests were performed using Prism software (GraphPad) using 5 mice per group.

**Table 1.**

| Target | Host species | Dilution | Catalog # | Company/origin |
| --- | --- | --- | --- | --- |
| *Antibodies used for immunohistochemistry* |  |  |  |  |
| GABAAR γ2 | Guinea pig | 1:2000 | - | Home-made |
| GABAAR α5 | Guinea pig | 1:3000 | - | Home-made |
| Gad-65/67 | Rabbit | 1:1000 | GC 3008 | Affiniti |
| Vglut1 | Guinea pig | 1:1000 | 135 304 | Synaptic Systems |
| PSD95 | Mouse | 1:1000 | 73-028 | NeuroMap |
| GFAP | Mouse | 1:2000 | MAB360 | Millipore |
| Iba-1 | Rabbit | 1:1000 | 019-19741 | Wako |
| Cd11b | Rat | 1:50 | 550282 | BD-Pharmingen |
| mBDNF N-9 | Mouse | 1:100 | BDNF #9 | DSHB |
| p75<sup>NTR</sup> | Rabbit | 1:200 | AB-NO1 | Advanced Targeting System |
| pro-BDNF #B240 | Mouse | 1:200 |  | Philip Barker |
| *Antibodies used for Western Blot* |  |  |  |  |
| GABAAR α1 | Rabbit | 1:600 | - | Home-made |
| GABAAR α2 | Rabbit | 1:1000 | - | Home-made |
| Gephyrin | Mouse | 1:1000 | <u>147 111</u> | Synaptic Systems |
| Gephyrin PS268 | Rabbit | 1:500 | - | Home-made |
| Gephyrin PS270 | Rabbit | 1:500 | - | Home-made |
| GSK3β | Rabbit | 1:1000 | ab124661 | Abcam |
| Erk1/2 | Rabbit | 1:4000 | 9102S | Cell Signaling |
| Phospho-Erk1/2<br>(Thr202/Tyr204) | Rabbit | 1:4000 | 9101 | Cell Signaling |
| BDNF | Rabbit | 1:3000 | Ab108319 | Abcam |
| Actin | Mouse | 1:10,000 | MAB1501 | Millipore |

### Microglia analysis

Iba1 staining was acquired using a spinning-disk confocal microscope (Nikon Ti2 coupled to Yokogawa CSU-W1 confocal scanning unit) with a Omicron modified Light HUB+ laser emitting at 488 nm and a CFI Plan Apochromat Lambda 60X oil objective (N.A 1.40, W.D. 0.13mm). 3D stack images of 25.5 µm were acquired with a z-step of 0.3 µm in layer II/III of the motor cortex (2-3 slices per animal, 5 animals per group). For the microglia density analysis, individual microglial cells were counted in FIJI, through colocalization of the Iba1 and DAPI stainings in 3D and the «Analyze Particles» function. The mean intensity and area covered by the Iba1 signal were analyzed on the maximum intensity projection of the stack in FIJI.

Imaris Software (Bitplane, Switzerland) was used for reconstruction of MCAO BDNF^wt/wt^ / CX3CR1^CreERT2+/-^ and BDNF^flox/flox^/ CX3CR1^CreERT2+/-^ cells. 3D rendering of microglial volume was based on Iba1 immunoreactivity, applying recorded algorithms with fixed thresholds for Iba1 signal intensity. Morphometric values were extracted per individual cells (BDNF^wt/wt^ / CX3CR1^CreERT2+/-^; Contra, n=68; Ipsi, n=72; BDNF^flox/flox^ / CX3CR1^CreERT2+/-^; Contra, n=82; Ipsi, n=94).

### Dendrite Reconstructions, Spine Quantification, cluster Quantification

3D confocal stacks were deconvolved with Huygens Essentials software (Scientific Volume Imaging, Hilversum, The Netherlands) using a full maximum likelihood extrapolation algorithm. Stacks were then imported and rendered using the Surpass function in Imaris software (Bitplane AG). Experimenter was blinded to conditions and treatment groups, spines were manually counted and using the ratio of the diameter and length of the head and neck of spines it was possible to distinguish between stubby, mushroom, and thin subtypes of dendritic spines. These classifications were based on previously established criteria ^34^. Lastly, *n* values for spine analysis represent ∼75-100 µm of dendrite from 1-2 cells imaged from each slice. The number and volume of gephyrin clusters were quantified using the Spot function of Imaris software, which differentiates puncta based on the fluorescence intensity.

### Electrophysiological Recordings and Analysis

Slices were transferred into a temperature-controlled chamber (25°C) mounted on an upright microscope (DM LFSA, Leica Microsystems) and continuously perfused with external solution. Patch recording electrodes were pulled from borosilicate glass (GC150TC; Clark Instruments, Old Sarum, Salisbury UK). All electrophysiological recordings were made using an Axopatch 200A amplifier (Molecular Devices, Sunnyvale, CA, U.S.A.).

AMPA-mediated mEPSCs were gathered from whole-cell voltage-clamp recordings of CA1 pyramidal neurons obtained at 25°C using electrodes with resistances of 4-5 MΩ and filled with intracellular solution containing (in mM): K-Gluconate, 120; EGTA, 1; HEPES, 10; Mg-ATP, 5; Na-GTP 0.5; NaCl, 5; KCl, 5; phosphocreatine, 10; 295 mOsm; pH adjusted with KOH to 7.3. mEPSCs were recorded at -60 mV and in the presence of 1 µM tetrodotoxin (TTX), 15 µM 3-[(R)-2-carboxypiperazin-4-yl]-propyl-1-phosphonic acid (CPP), 100 µM picrotoxin, and 1 µM CGP55845 in the external Tyrode’s solution. Access resistance was monitored with brief test pulses at regular intervals (2-3 min) throughout the experiment. Access resistance was usually 10-13 MΩ and data were discarded if the resistance deviated more than 10% through the course of the experiment. Series resistance of the access pulse and decay time was also used for the calculation of total membrane capacitance. After the holding current had stabilized, data were recorded at a sampling frequency of 10 kHz and filtered at 2 kHz for 10 to 15 min.

GABA_A_R-mediated mIPSCs were gathered from whole-cell voltage-clamp recordings of CA1 pyramidal neurons obtained at 25°C using electrodes with resistances of 4-5 MΩ and filled with intracellular solution containing (in mM): CsCl, 140; NaCl, 4; 0.5, CaCl2; HEPES, 10; EGTA, 5; QX-314, 2; Mg-ATP, 2; Na-GTP 0.5; 290 mOsm; pH adjusted with CsOH to 7.36. mIPSCs were recorded at -60 mV and in the presence of 1 µM TTX, 25 µM CPP, 5 µM CGP55845, 5 µm 6-cyano-7-nitroquinoxaline-2,3-dione (CNQX) and 0.3 µm strychnine in external Tyrode’s solution. Access resistance was monitored with brief test pulses at regular intervals (2-3 min) throughout the experiment. After the holding current had stabilized, data were recorded at a sampling frequency of 10 kHz and filtered at 2 kHz for 10 to 15 min.

All mEPSCs and mIPSCs were detected offline using the Mini Analysis Software (Synaptosoft, Decatur, GA, USA). The amplitude threshold for mEPSC and mIPSCs detection was set at four times the root-mean-square value of a visually event-free recording period. From every experiment, 5 min of stable recording was randomly selected for blinded analysis of amplitude and inter-event interval. The data obtained was then used to plot cumulative histograms with an equal contribution from every cell. For statistical analysis, data were averaged for every single cell. It should be noted that the amplitude analysis was conducted only on single mEPSCs and mIPSCs that did not have subsequent events occurring during their rising and decaying phases. For frequency analysis, all selected events were considered.

### Pharmacological Treatments

To scavenge BDNF, slices were treated with TrkB-Fc (R&D Systems; Minneapolis, MN, USA), a fusion protein in which the BDNF binding site of the TrkB receptor replaces the Fc fragment of a human IgG1 antibody. We found that TrkB-Fc treatment to hippocampal cultures for 24 h downregulated TrkB receptor phosphorylation (data not shown). TrkB-Fc was diluted in culture media at a final concentration of 10 µg/ml and treatment began immediately following induction of OGD. ERK activation was inhibited using 30 µM of MEK inhibitor PD98059 (Tocris Biosciences, ON, Burlington, Canada), GSK3β activity was inhibited using 25 µM GSK3β-IX (Tocris Biosciences). PD98059 and GSK3β-IX were diluted in dimethyl sulfoxide (Invitrogen) and treatment began overnight prior to OGD induction, removed during induction and continued for 90 min after. Control sister cultures were treated with control culture media containing dimethyl sulfoxide only. The following function blocking antibodies were used: 1:200 anti-p75^NTR^ (kind gift from Dr. P. Barker, McGill University, Montreal, QC, Canada; Rex antibody for more information see ^78^, 1:200 anti-proBDNF (kind gift from Dr. Philip Barker, McGill University, Montreal, Canada) and 1:100 anti-mBDNF (N-9, Developmental Studies Hybridoma Bank, University of Iowa, IA, USA; for more information see ^79^. Function blocking antibody treatment began 2 h prior to OGD induction, removed during induction and continued for 90 min after.

### Biolistic Gene Transfection

Cartridges were prepared according to manufacturer’s protocol (Bio-Rad, Helios Gene Gun). Briefly, 15 mg of gold particles (1 µm diameter) were first coated with 0.05 M spermidine. 15 µg of plasmid DNA expressing tdTomato and 45 µg of wildtype gephyrin-GFP (gephyrin_WT_-GFP) or dephosphorylation mutant gephyrin-GFP S268A/S270A (gephyrin_S268A/S270A_-GFP). Plasmids were then precipitated onto the particles by adding CaCl2. The coated particles were resuspended into 100% ethanol and infused into Tefzel tubing, which were then coated with the particles. Coated tubing was cut into 0.5 inch cartridges which were then transfected into mature organotypic slice cultures by shooting at a distance of 2 cm with a pressure of 200 psi through a nylon mesh. Following 48 h, slices which expressed target plasmids in CA1 pyramidal neurons were processed with OGD or control Tyrode solution.

### Western blotting

24 h following MCAO, mice were killed by decapitation and brains were dissected on ice. The ipsi- and contralateral cortices were immediately transferred to lysis buffer (50 mM Tris, pH 7.6, 150 mM NaCl, 1% Triton X-100, CompleteMini Protease Inhibitor Mixture, Roche). The cortices were homogenized and incubated on ice for 1 hour. Lysates were centrifuged at 20,000 RPM for 30 min at 4°C, and supernatants were stored at -80°C. Samples were run on Tris-glycine polyacrylamide gels and proteins were transferred to PVDF membranes. Primary antibodies (see Table 1) were incubated in Tris-buffered saline with 0.05% Tween 20 (TBST), including 5% WesternBlocking Solution (Roche) overnight at 4°C. Membranes were washed 5 x 5min in TBST. HRP-coupled donkey secondary antibodies (1:30,000) and fluorescent-coupled donkey secondary antibodies (1:20,000) were incubated for 30 min at room temperature, and membranes were washed again 5 times 5 min in TBST. Fluorescent signals were captured using the Odyssey® CLx Imager. SuperSignal West Pico ChemiluminescentSubstrate (Thermo Fisher Scientific) was applied to visualize HRP labelled antibodies and developed using the FUJIFILM Luminescent Image Analyzer LAS-1000 plus & Intelligent Dark Box II (Fujifilm). Images were analyzed using ImageJ and statistical tests were performed using Prism software (GraphPad) using a minimum of 4 mice per group. Western blot membrane stripping for restaining was performed for p-ERK1/2 and ERK1/2 antibodies using a mild stripping protocol from abcam. Briefly, membranes were incubated twice for 5-10 min with mild stripping buffer (200mM gylcine, 20mM SDS, 0.01% Tween 20, pH 2.2) followed by 2 x 10min incubation with PBS and 2 x 5min incubations with TBST. Efficiency of stripping was checked by incubating with chemiluniscent detection. When stripping was judged satisfactory, the membranes were rinsed and incubated with primary antibody.

### Statistical Analysis

An a priori power analysis was conducted using data reported in Figures 5-8 obtained with the MCAO surgery procedure. Combined with type-1 error set to 0.05. Power set to 0.8 (1-beta-error), we determined the effect size with a group size of 5. These indicated an inter individual variation (SD) of 10-15%. We used multiple group comparison test (Bonferroni) to make maximal number of comparisons between groups.

For the slice culture comparisons we performed 2-way ANOVA and independent t-tests on the following figures: Figure 2b, d, e, f’, f, g, g’; Figure 3b, c, d, f, g, h; Figure 4b, c, d, g, h, i. The t-tests were performed to confirm significance between conditions as 2-way ANOVA and post hoc test does not cover all the comparisons of our interest. These t-test p values are unprotected, therefore we adjusted them using Bonferroni correction (e.g. if the set of the data is compared 5 times, we need to multiply the p-value by 5). The listed significant comparisons all have protected p-values by Bonferroni post-test. *** p<0.001; **p<0.01; *p<0.05.

Comparison between two groups were made using two-tailed independent Student’s *t*-test. Comparisons between multiple groups and treatments were made using two-way ANOVA with *post hoc* Bonferroni multiple comparison test. Comparisons between multiple groups were made using one-way ANOVA. Cumulative probability plots were compared using Kolmogorov-Smirnov test for probability distributions. Results are expressed as mean ± sd.

## Author Contributions

SKT and RAM designed the study. ZST and PKYC performed and analyzed the OGD electrophysiology. RG performed and analysed the morphology data from OGD experiments. ZST, PKYC and RG supported data collection and analysis. MV performed the MCAO, TC, MV, FM, RP and SS performed and analysed the mouse MCAO data. DC wrote the macro for morphology analysis. SBN performed data analysis. RCP, PAB, SAB and JK provided technical expertise, infrastructure support and biological reagents for this study. RG, PKYC, ZST, FM prepared OGD figures, TC, SS, SBN, MV, FM and SKT prepared MCAO figures. All authors contributed to writing and editing the manuscript.

## Acknowledgments

We thank Dr. J-M. Fritschy for antibody reagents and feedback on the manuscript; F. Charron for preparation and maintenance of organotypic slice cultures; G.Bosshard for preparation of microglia cultures; Plexxikon for sharing their PLX5622 chow formulated compound for our studies. This project was initiated as part of ZNZ-McGill-Oxford tripartite collaboration. This work was supported by grants from CIHR to RAM (grant ID: MOP 133611), Savoy Foundation to RAM, Swiss National Science Foundation (320030_179277), ERA-NET NEURON (32NE30_173678/1), the Synapsis Foundation and the Vontobel foundation to JK, Swiss National Science Foundation grants to SKT (31003A_159867, 310030_192522), internal UZH funding to SKT, Forschungskredit Candoc grant to TC and Forschungskredit postdoc grant to MV.

---

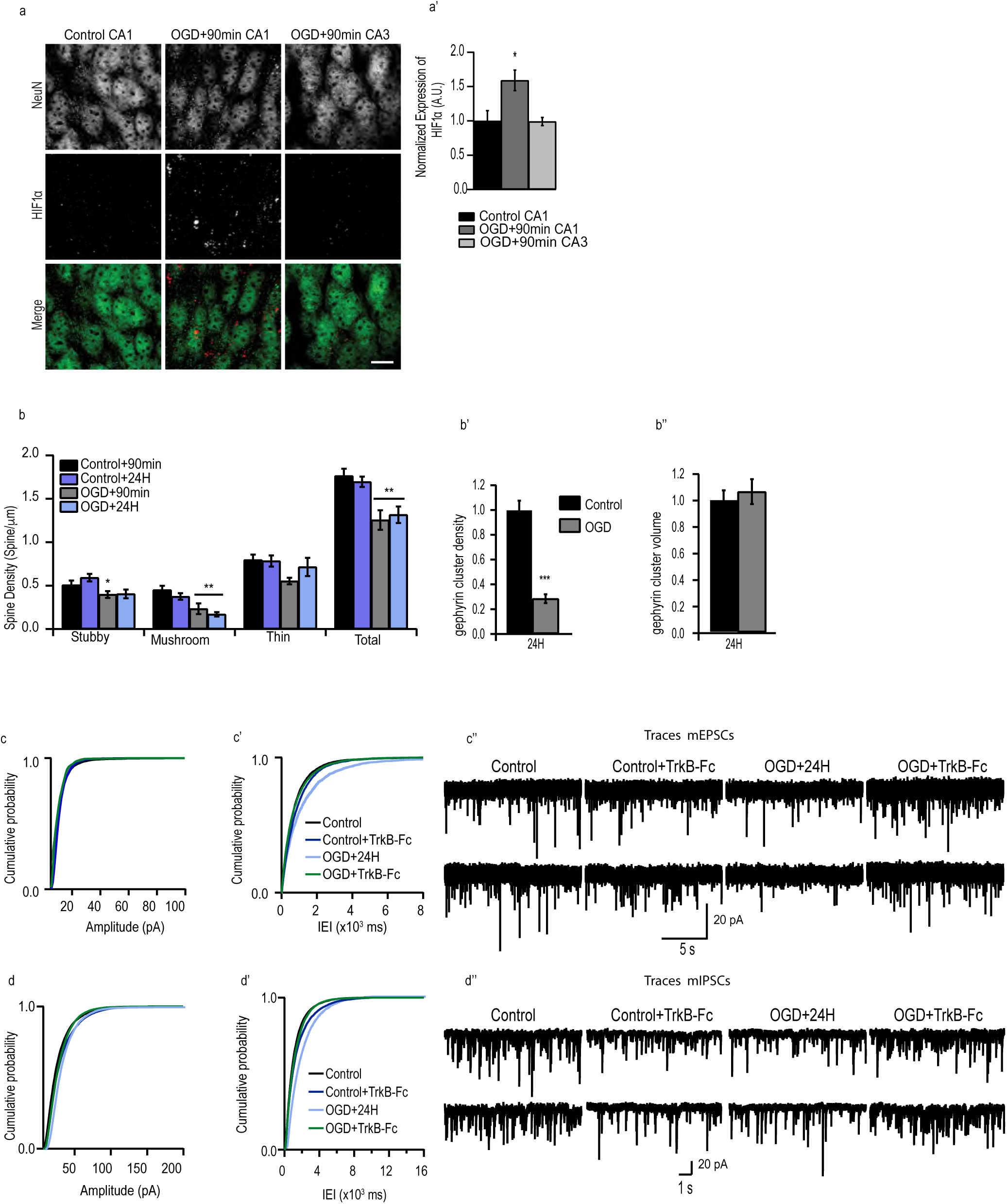

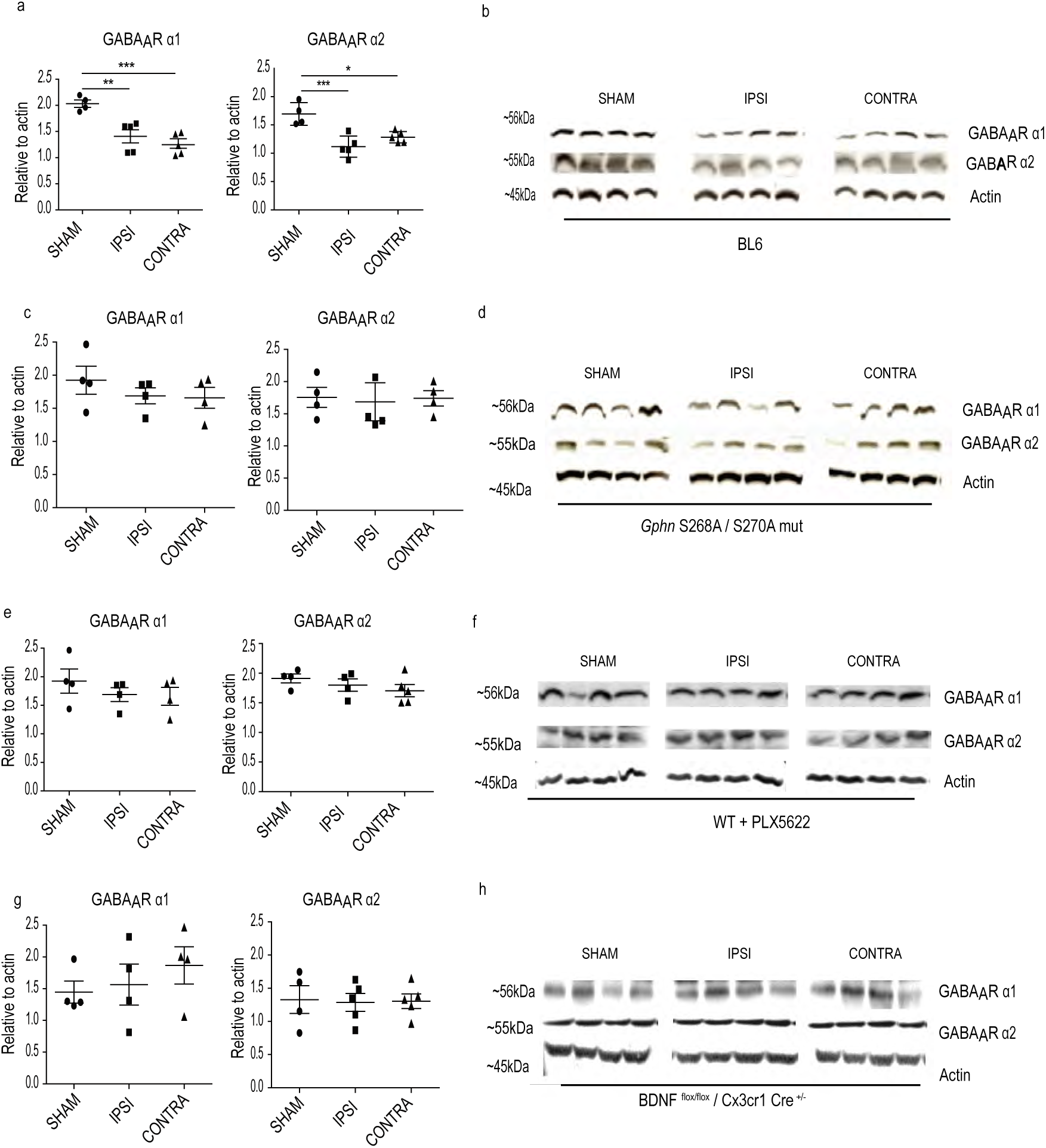

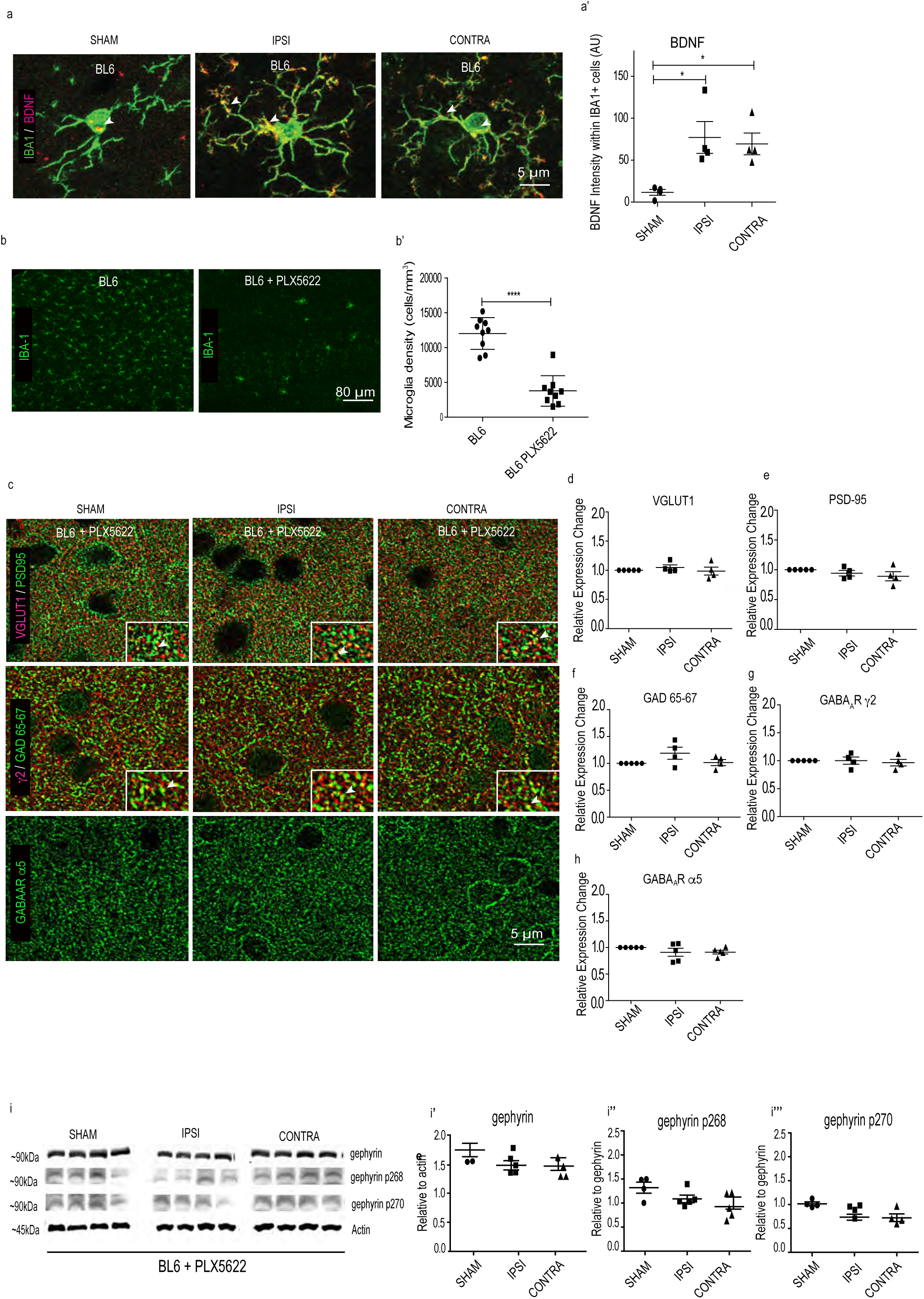

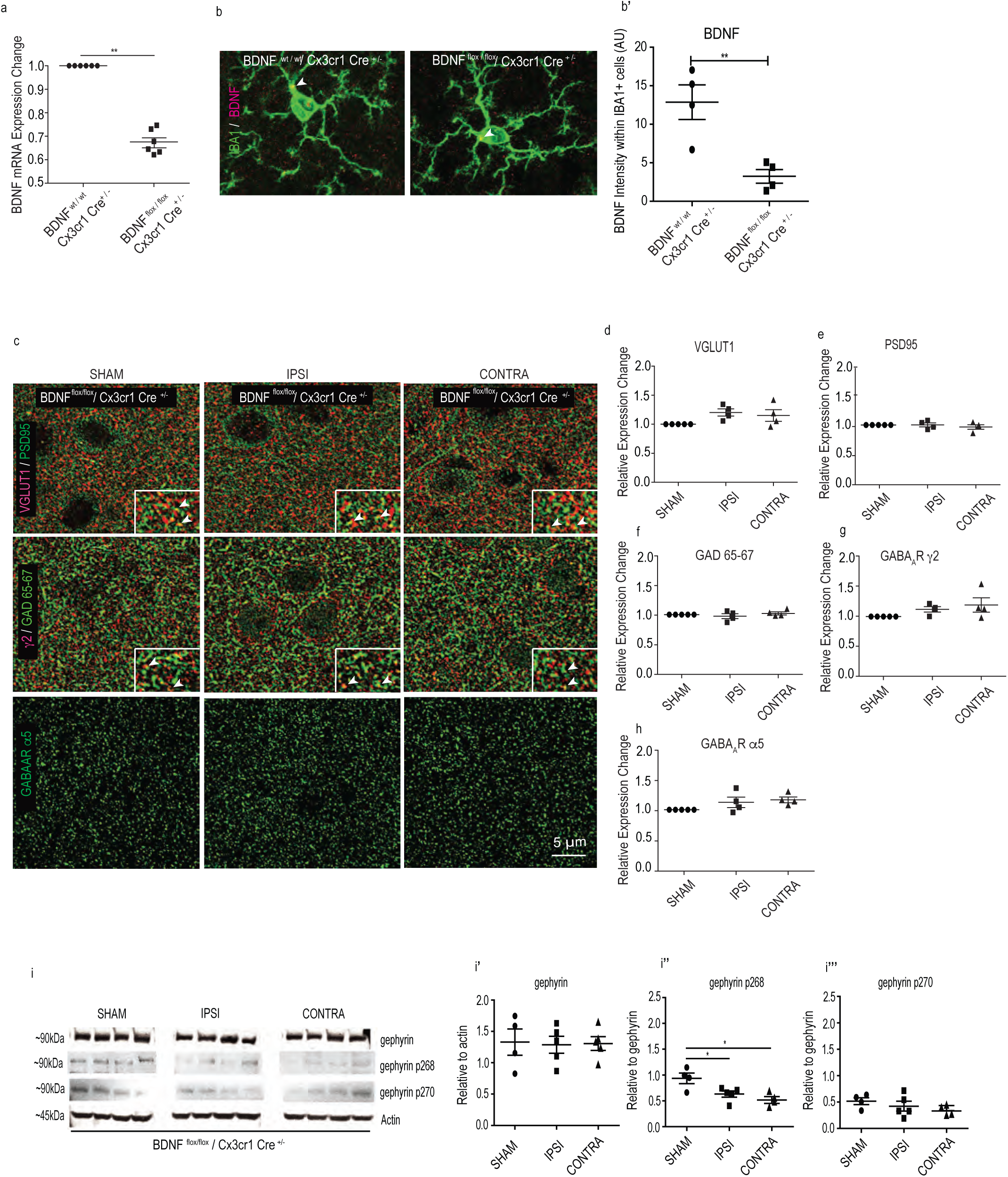

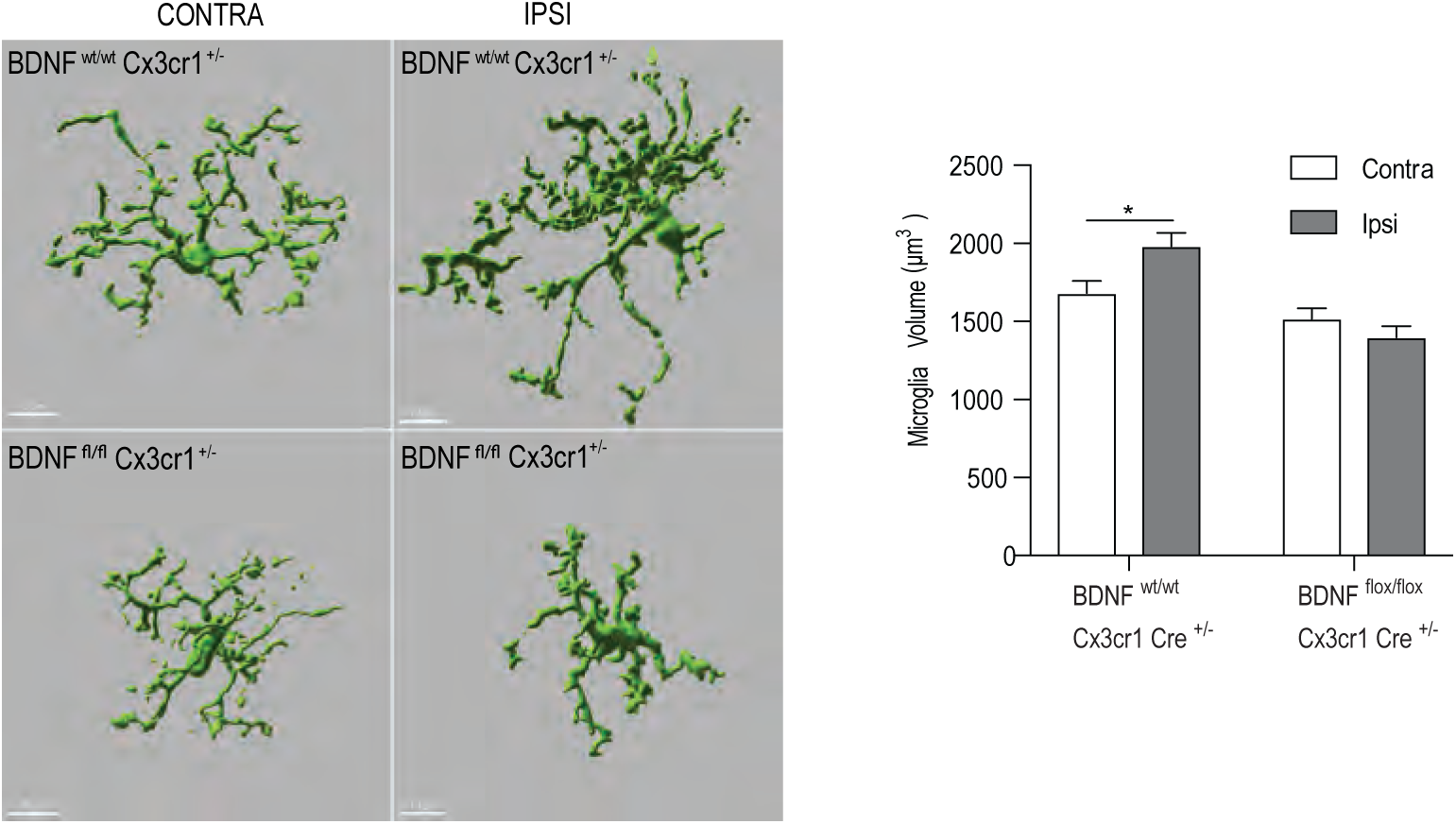

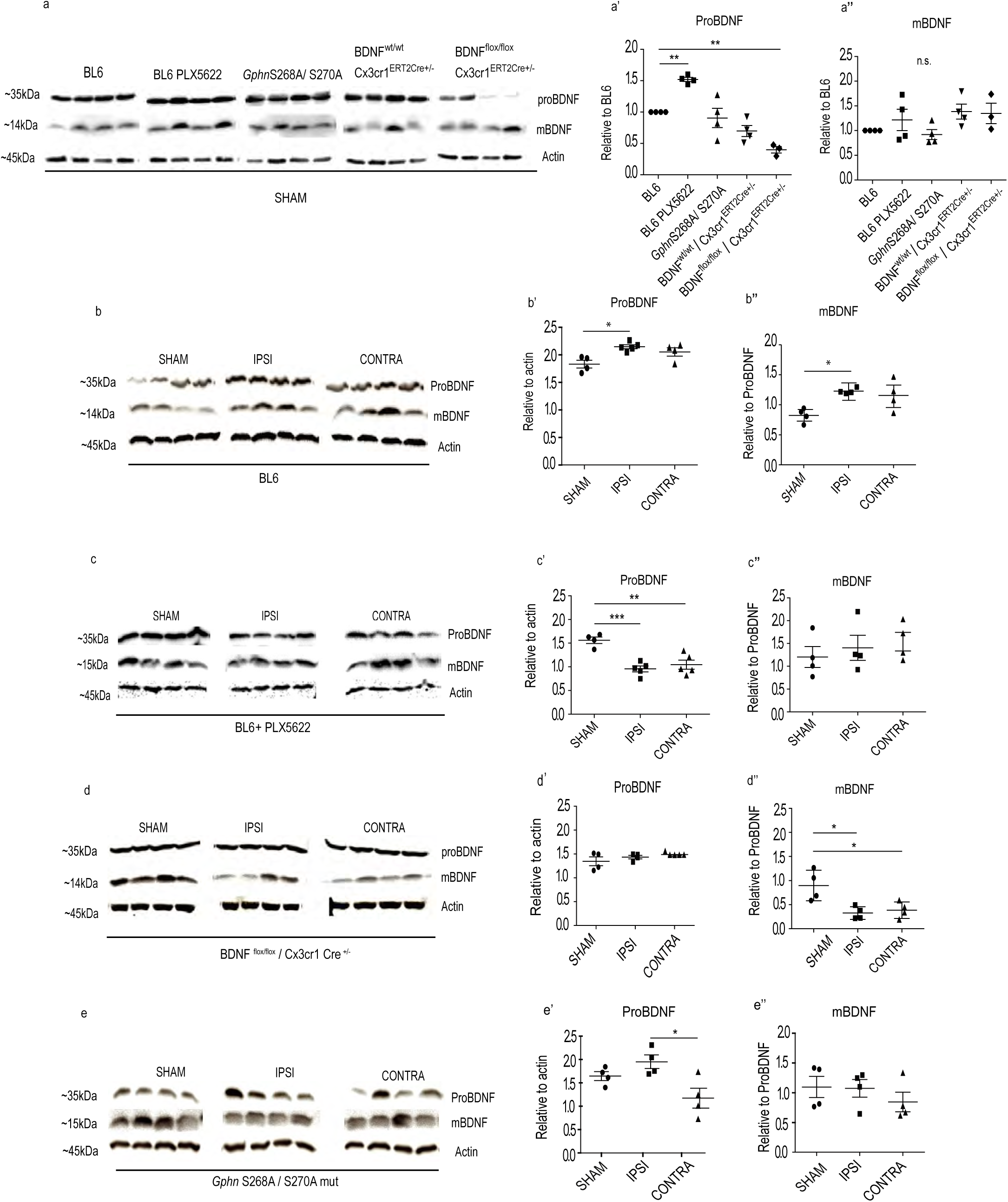

