## Supplementary material for "Blocking gephyrin phosphorylation or microglia BDNF signaling prevents synapse loss and reduces infarct volume after ischemia": Suppl. Figure legends

*1. Institute of Pharmacology and Toxicology, University of Zurich, Winterthurerstrasse 190, CH 8057, Switzerland*
*2. Department of Pharmacology and Therapeutics, McGill University, 3649 Prom. Sir-William-Osler, Montreal, QC, H3G 0B1, Canada*
*3. Department of Biology, University of British Columbia, 3187 University Way, ASC 413 Kelowna, BC, V1V 1V7, Canada*
*4. Department of Biomedical Sciences, University of Lausanne, Rue du Bugnon 7 - CH-1005 Lausanne*
*5. Institute for Biomedical Engineering, University of Zurich and ETH Zurich, Wolfgang-Pauli-Strasse 27, CH 8093, Switzerland*
* equal contribution

***Suppl. Figure 1 – OGD causes glutamatergic and GABAergic synapse loss via BDNF signaling.*** **(a)** HIF1α expression in CA1 pyramidal neurons at 90min after OGD in hippocampal slice culture. **(a')** Quantification of HIF1α expression in CA1 pyramidal neurons. **(b)** Quantification of dendritic spines at 90 min and 24 h after OGD. The spines were categorized into stubby, mushroom and long thin subtypes (*$p < 0.05$ and **$p < 0.01$, two-tailed independent Student's t-test). Total spine density (spines/µm of dendrite). **(b')** Quantification of gephyrin cluster density 24 h after OGD (consisting of five 512x512 pixel z-planes each; ***$p < 0.0001$, two-tailed independent Student's t-test). **(b'')** Quantification of the total gephyrin cluster volume at 24 h after OGD (***$p < 0.0001$, two-tailed independent Student's t-test). **(c)** Cumulative probability histogram of mEPSC amplitude in TrkB-Fc treated neurons at 24 h post OGD ($p < 0.05$ Kolmogorov-Smirnov test). **(c')** Cumulative probability histogram of mEPSC IEIs ($p < 0.05$, Kolmogorov-Smirnov test). **(c'')** Example traces of AMPA-mediated mEPSC recordings after TrkB-Fc treatment and OGD. **(d)** Cumulative probability histogram of mIPSC amplitudes in TrkB-Fc treated neurons at 24 h post OGD ($p < 0.05$, Kolmogorov-Smirnov test). **(d')** Cumulative probability histogram of mIPSC IEIs (p <

0.05, Kolmogorov-Smirnov test). **(d'')** Example trace of mIPSCs after TrkB-Fc treatment and OGD .

***Suppl. Figure 2 – a1 and a2 GABA$_A$R expression levels in different mouse lines after OGD.*** **(a-b)** WB analysis and quantification for α1 and α2 GABA$_A$R in BL6 WT mice at 24 h after MCAO. **(c-d)** WB analysis and quantification for α1 and α2 GABA$_A$R in *Gphn*S268A/S270A mutant mice at 24 h after MCAO. **(e-f)** WB analysis and quantification for α1 and α2 GABA$_A$R in BL6 WT mice treated with PLX5622 at 24 h after MCAO. **(g-h)** WB analysis and quantification for α1 and α2 GABA$_A$R in BDNF$^{flox/flox}$ / Cx3Cr1$^{CreERT2+/-}$ mice at 24 h after MCAO. Quantification expressed as mean ± s.d. (N=4); *P<0.05; **P<0.01; ***P<=0.001 (One-way ANOVA, Bonferroni multiple comparison test).

***Suppl. Figure 3 – Microglia depletion stabilize synapse changes after OGD*** **(a)** BDNF expression within microglia in BL6 WT sham and MCAO animals. **(a')** Quantification of BDNF intesity within IBA1 positive cells. **(b)** Example images from BL6 WT mice treated with PLX5622 sham and MCAO. **(b')** Microglia depletion after PLX5622 chow treatment for 7 days (1200mg/kg of chow). **(c)** Example images from BL6 WT PLX5622 treated MCAO animals, quantified for **(d-e)** VGLUT1, PSD95 and **(f-h)** GAD65-67, GABA$_A$R γ2 and GABA$_A$R α5 expression change relative to sham in BL6 WT PLX5622 treated MCAO animals. **(i)** WB analysis for total gephyrin, phospho gephyrin-S268 and phospho gephyrin-S270 levels in BL6 WT mice treated with PLX5622 at 24 h after MCAO. **(i'-i''')** Quantification of WB for total gephyrin, phospho gephyrin-S268 and phospho gephyrin-S270 levels in BL6 WT mice treated with PLX5622 at 24 h after MCAO. Quantification expressed as mean ± s.d. (n=5); p>0.05 (One-way ANOVA, Bonferroni multiple comparison test).

***Suppl. Figure 4 – Bdnf deletion from microglia stabilize synapses after MCAO.*** **(a)** Quantitative RT-PCR for *Bdnf* mRNA changes in cultured microglia from $BDNF^{wt/wt}$ / $CX3CR1^{CreERT2+/-}$ and $BDNF^{flox/flox}$ / $CX3CR1^{CreERT2+/-}$ mice. **(b)** Example images of BDNF protein expression within microglia from $BDNF^{wt/wt}$ / $CX3CR1^{CreERT2+/-}$ and $BDNF^{flox/flox}$ / $CX3CR1^{CreERT2+/-}$ mice. **(b')** Quantification of BDNF intensity within IBAI cells in $BDNF^{wt/wt}$ / $CX3CR1^{CreERT2+/-}$ and $BDNF^{flox/flox}$ / $CX3CR1^{CreERT2+/-}$ mice. **(c)** Example images from $BDNF^{flox/flox}$ / $CX3CR1^{CreERT2+/-}$ sham and MCAO mice showing VGLUT1/PSD95 (column 1), $GABA_AR$ $\gamma2$/ GAD65-67 (column 2), $GABA_AR$ $\alpha5$ (column 3). **(d-e)** Quantification of VGLUT1, PSD95 expression change in MCAO animals relative to sham. **(f-h)** Quantification of GAD65-67, $GABA_AR$ $\gamma2$ and $GABA_AR$ $\alpha5$ expression change in MCAO animals compared to sham. **(i)** WB analysis for total gephyrin, phospho-gephyrin S268 and S270 in $BDNF^{flox/flox}$ / $CX3CR1^{CreERT2+/-}$ sham and MCAO mice. **(i'-i''')** Quantification of total gephyrin, phospho gephyrin S268 and S270. Morphology analysis shown as mean ± s.d. (n=4 animals); P=0.55 (Two-way ANOVA, Bonferroni multiple comparison test; F=(2,9)=8.7).

***Suppl. Figure 5 – Microglia volume change in $BDNF^{flox/flox}$ / $CX3CR1^{CreERT2+/-}$ mice after MCAO.*** **(a)** Example image of 3D volumetric analysis of IBA1 positive cells in $BDNF^{wt/wt}$ / $CX3CR1^{CreERT2+/-}$ mice and $BDNF^{flox/flox}$ / $CX3CR1^{CreERT2+/-}$ mice ipsi- and contra-lateral sides. (b) Quantification of volume analysis in $BDNF^{wt/wt}$ / $CX3CR1^{CreERT2+/-}$ mice and $BDNF^{flox/flox}$ / $CX3CR1^{CreERT2+/-}$ mice ipsi- and contra-lateral sides. Data shown as mean ± s.d. (n=5 animals); *P<0.05

***Suppl. Figure 6 –BDNF expression difference between mouse lines at baseline and after MCAO.*** **(a-a'')** WB analysis for proBDNF and mBDNF from parietal cortex L2/3 in BL6 WT mice, BL6 WT mice treated with PLX5622, *Gphn*S268A/S270A mutant mice, $BDNF^{wt/wt}$ /

Cx3cr1$^{ERT2Cre+/-}$ and BDNF$^{flox/flox}$ / Cx3cr1$^{ERT2Cre+/-}$. **(b)** WB analysis of cortex L2/3 in WT mice for changes in proBDNF and mBDNF levels 24 h following MCAO. **(b'-b'')** Quantification for proBDNF (One-way ANOVA, Bonferroni multiple comparison test; F(2,10)=7.3; P=0.01) and mBDNF (One-way ANOVA, Bonferroni multiple comparison test; F(2,9)=6.8; P=0.015) from cortex of ipsi- and corresponding contra-lateral hemispheres in WT mice 24 h post MCAO. **(c)** WB analysis of cortex L2/3 in BL6 WT mice treated with PLX5622 for changes in proBDNF and mBDNF levels 24 h post MCAO. **(c'-c'')** Quantification for proBDNF (One-way ANOVA, Brown-Forsythe test; F(2,11)=15.8; P=0.0006) and mBDNF from ipsi- and corresponding contra-lateral hemispheres in BL6 WT mice treated with PLX5622. **(d)** WB analysis of cortex L2/3 in BDNF$^{flox/flox}$ / CX3CR1$^{CreERT2+/-}$ for proBDNF and mBDNF levels 24 h following MCAO. **(d'-d'')** Quantification for proBDNF and mBDNF (One-way ANOVA, Brown-Forsythe test; F(2,9)=8.044; P=0.0099) from ipsi- and corresponding contra-lateral hemispheres in BDNF$^{flox/flox}$ / CX3CR1$^{CreERT2+/-}$ mice. **(e)** WB analysis of cortex L2/3 in *Gphn*S268A/S270A mutant mice for proBDNF (One-way ANOVA, Brown-Forsythe test; F(2,9)=6.1; P=0.02) and mBDNF levels 24 h following MCAO. **(e'-e'')** Quantification for proBDNF and mBDNF from ipsi- and corresponding contra-lateral hemispheres in *Gphn*S268A/S270A mutant mice.
